# Single Nucleus Multiome Analysis Reveals Early Inflammatory Response to High-Fat Diet in Mouse Pancreatic Islets

**DOI:** 10.1101/2025.04.01.646568

**Authors:** Isabell Victoria Strandby Ernst, Lasse Lehtonen, Sille Marie Nilsson, Freja Louise Nielsen, Ann-Britt Marcher, Susanne Mandrup, Jesper Grud Skat Madsen

## Abstract

In periods of sustained hyper-nutrition, pancreatic β-cells undergo functional compensation through transcriptional upregulation of gene programs driving insulin secretion. This adaptation is essential for maintaining systemic glucose homeostasis and metabolic health. Using single nuclei multiomics, we have mapped the early transcriptional compensation mechanisms in murine islets of Langerhans exposed to high-fat diet (HFD) for one and three weeks. We show that β-cells exhibit the largest transcriptional response to HFD, characterized by early activation of proinflammatory eRegulons and downregulation of β-cell identity genes, particularly in a distinct subset of β-cells. Our observations translate to humans, as we observe an increase in the inflammatory gene signatures in human β-cells in pre-diabetes and diabetes. Collectively, these observations point to cellular cross-talk through proinflammatory signaling as a central and early driver of β-cell dysfunction that limits the compensatory capacity of β-cells, which is closely linked to the development of diabetes.

## Introduction

The increasing incidence of type 2 diabetes mellitus (T2DM), which coincides with a growing prevalence of obesity and a sedentary lifestyle^1,2^, constitutes a major global health threat. T2DM is a pleiotropic disease, which involves both decreased responsiveness to insulin in multiple tissues and inadequate insulin secretion by pancreatic β-cells. Thus, the ability of the β-cells to adapt to increased nutrient intake and secrete adequate amounts of insulin plays a key role in preventing the development of T2DM.

Functional compensation to meet the increased demand for insulin involves both increased β-cell mass and upregulation of insulin secretory capacity. Cellular adaptation through transcriptional and metabolic regulation plays a crucial role during functional compensation, where β-cells become more sensitive to glucose, exhibit enhanced glucose uptake and metabolism, and upregulate pathways involved in the production and secretion of insulin^3^. Additionally, excess nutrients, such as glucose and free fatty acids are directed away from harmful metabolic processes through storage, conversion, or export, which enables the β-cell to maintain function^4^. The loss of the ability to compensate represents a crucial tipping point in the development of T2DM, and understanding the underlying mechanisms is essential to understanding T2DM.

Single-cell technologies have shed light on the functional and transcriptional heterogeneity of β-cells. For example, several single-cell RNA-seq (scRNA-seq) studies have observed heterogeneity in the functional maturity of β-cells in islets from adult humans and mice^5–7^. Specifically, it has been suggested that adult human β-cells exist in at least two or three different states^6,8–13^ which differ in terms of their epigenome, transcriptome, and electrophysiology. Studies have observed a shift in the relative abundance of β-cell subpopulations. One study, using machine learning on chromatin accessibility data from nondiabetic, pre-T2DM, and T2DM donors, identified two transcriptionally and functionally distinct β-cell sub-types that shift in abundance during T2DM progression^8^. Another study found that the distribution of four antigenically identified β-cell subtypes, all present in nondiabetic individuals, was altered in T2DM individuals^10^, suggesting that control of β-cell function is at least partly regulated through cell state transition.

To gain a deeper understanding of the dynamic gene regulatory mechanisms controlling the cell state and function of islet cells during functional compensation, we fed mice a high-fat diet (HFD) for one and three weeks to induce a compensatory response. We used single-nucleus multi-omics to simultaneously analyze changes in chromatin accessibility using single-nucleus assay for transposase-accessible chromatin using sequencing (snATAC-seq) and gene expression using single-nucleus RNA-seq (snRNA-seq) in islets of Langerhans. Here, we show that of the different cell types in the islets, β-cells are most strongly affected by short-term HFD. We map gene regulatory networks and discover a subpopulation of β-cells that are marked by proinflammatory gene signatures that correlate to systemic metabolic traits in mice. This gene signature was also detected in β-cells from humans who had prediabetes or diabetes, suggesting that inflammation may be an important regulator of the capacity for functional compensation in β-cells.

## Results

### Characterization of Islets of Langerhans during Short-Term High Fat Diet

To gain cell type-resolved insight into the transcriptional events regulating compensation in islets of Langerhans, we initially established and characterized an experimental setup using short-term HFD feeding in mice to induce a compensatory response. Eight-weeks-old male C57BL/6JBomTac mice were first exposed to low fat diet (LFD) for three weeks, whereafter they were either maintained on the LFD (control) or switched to HFD for one or three weeks (**Figure 1A, Table S1**). HFD-fed mice exhibited increased body weight gain, fasting hyperglycemia, hyperinsulinemia, mild glucose intolerance, and onset of insulin resistance compared to LFD controls already after one week (**Figure 1B-I, Figure S1A-B, Table S2-6**). These phenotypic differences between HFD and LFD-fed mice were similar after three weeks of HFD, except for a more pronounced increase in body weight. These results are consistent with previous studies^14–18^ and indicate that the one and three weeks-time points are adequate for inducing a compensatory response. In line with these observations, the secretory capacity of pancreatic β-cells, estimated by the structure parameter inference approach (SPINA-Gβ), increased after one week of HFD compared to the LFD controls, although this difference did not reach statistical significance (p = 0.065). This suggests that after one week of HFD, the β-cell compensation has been induced.

**Figure 1:**
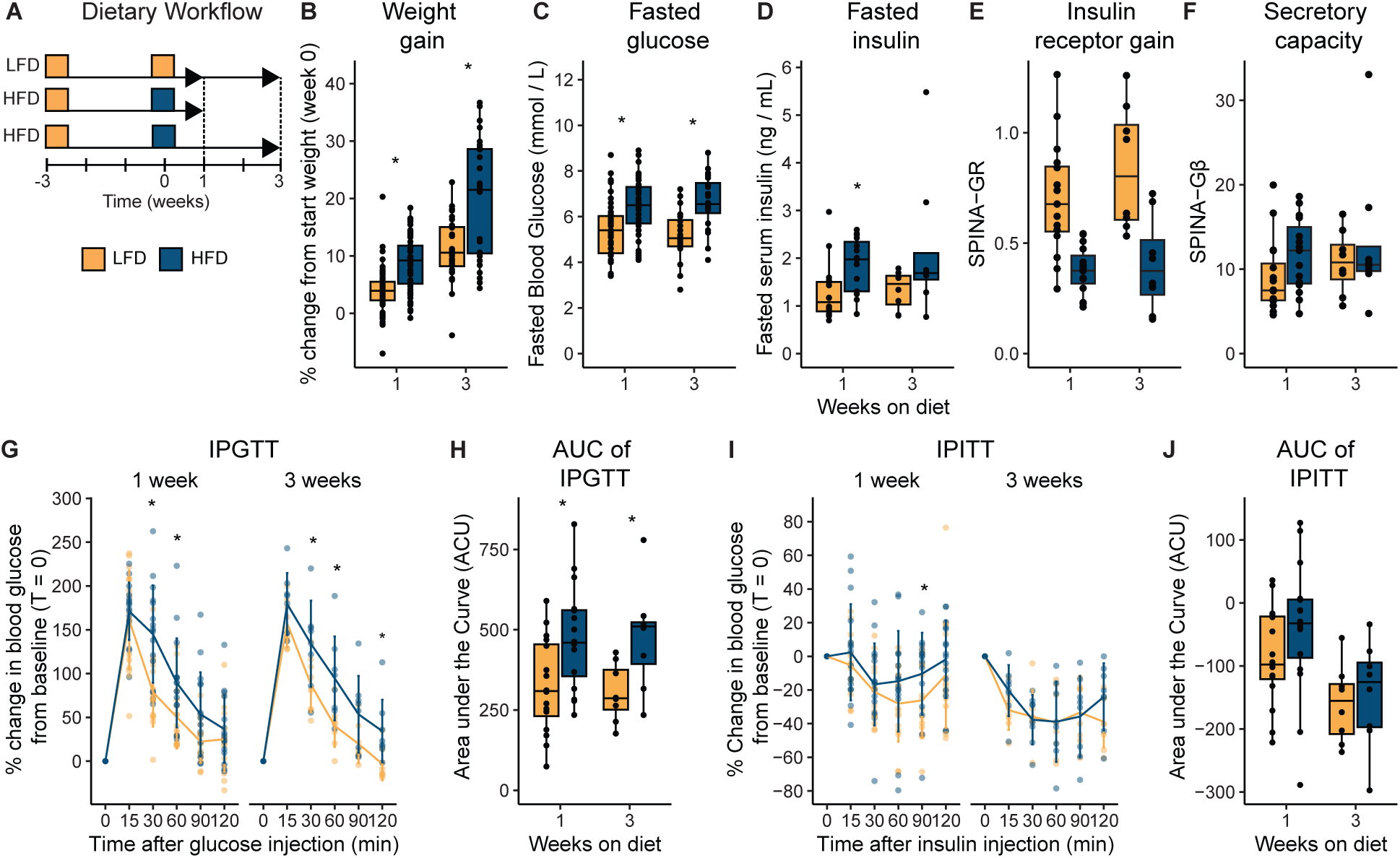
Short-term HFD leads to increased body weight gain, fasting hyperglycemia, hyperinsulinemia, mild glucose intolerance and onset of insulin resistance. **A)** Schematic overview of the experimental setup: 8-week-old male C57BL/6JBomTac mice underwent a three-week acclimatization period during which they were fed a low-fat diet (LFD). Subsequently, mice were divided into groups receiving either LFD (orange, as a control) or high-fat diet (blue, HFD) for one or three weeks. **B)** Weight gain measured as a percentage of starting weight at day 0 for each animal after one week (n = 66-70 per diet) and three weeks (n = 28 per diet). **C)** Fasting blood glucose levels in mmol/L at one week (n = 48 per diet) and three weeks (n = 24 per diet). **D)** Fasting serum insulin levels in ng/mL at one week (n = 15-16 per diet) and three weeks (n = 8 per diet). **E)** Structure parameter inference approach (SPINA)-Carb of insulin receptor gain / insulin sensitivity (SPINA-GR) at one week (n = 15-16 per diet) and three weeks (n = 8 per diet). **F)** Structure parameter inference approach (SPINA)-Carb of secretory capacity / β-cell function (SPINA-Gβ) at one week (n = 15-16 per diet) and three weeks (n = 8 per diet). **G)** Intraperitoneal glucose tolerance test (IPGTT) at one week (n = 16 per diet) and three weeks (n = 8 per diet). Data is shown as a percentage change from blood glucose at T = 0. **H)** The data points shown in the IPGTT (G) were used to calculate the area under the curve. (AUC). **I)** Intraperitoneal insulin tolerance test (IPITT) at one week (n = 16 per diet) and three weeks (n = 8 per diet). Data is shown as a percentage change from serum insulin levels at T = 0. **J)** The data points shown in the IPITT (I) were used to calculate the area under the curve.

To investigate the single cell-resolved transcriptional response of islets to HFD, we jointly measured both chromatin accessibility and gene expression using snATAC-seq and snRNA-seq, respectively, in single nuclei from isolated islets of Langerhans from mice fed with LFD for one (n = 2) or three weeks (n = 1), or HFD for one (n = 2) or three weeks (n = 2). Following rigorous quality filtering of both the snRNA-seq and snATAC-seq data (**Figure S2A-J, Table S7**), a total of 20,566 nuclei were included in the downstream analyses. We performed dimensional reduction and batch integration for each modality (**Figure S2K**). Using weighted nearest neighbor (WNN) analysis^19^, we created a joint embedding incorporating information from both data modalities and used it for unsupervised clustering (**Figure 2A, Figure S2L**). We merged and labeled the clusters as eight different cell types based on marker gene expression, predicted gene scores, and pseudo-bulk chromatin accessibility tracks of known major marker genes in each cluster (**Figure 2B-C, Figure S2M, Table S8**). Both data modalities contributed equally to identifying the endocrine cell type populations based on modality-specific weights (**Figure 2D**), except for γ-cells, which were masked in the snATAC-seq data (**Figure 2E**) but not in the snRNA-seq data (**Figure 2F**).

**Figure 2:**
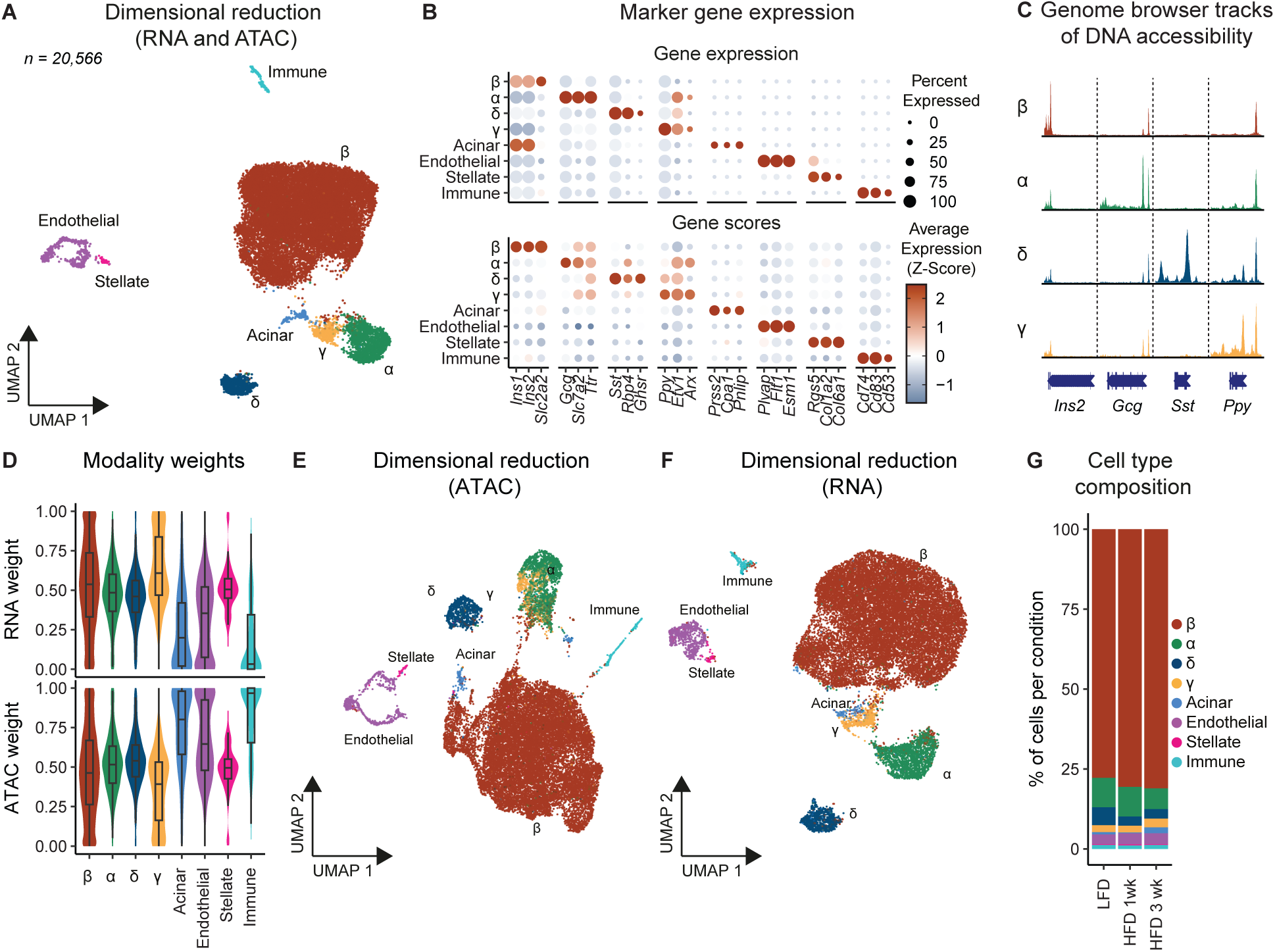
Classification of Islet Cell Types through Integrated snRNA and snATAC-seq Analyses. **A)** 11-week-old male C57BL/6JBomTac mice were fed either a low-fat diet (LFD) (n = 3) or high-fat diet (HFD) for one (1wk) (n = 2) and three (3wk) (n = 2) weeks. At the end of the experiment, the mice were sacrificed, and islets of Langerhans were isolated. Nuclei from islets were purified and used for Single Cell Multiome ATAC + Gene expression sequencing. The scatterplot shows Uniform Manifold Approximation and Projection (UMAP) visualization of clustering based on a weighted nearest neighbor (WNN) graph, based on the weighted average of RNA and ATAC similarities (n = 20,566). Data points are colored by cell type as indicated on the figure. **B)** Dot plot showing the percentage of cells expressing the gene (dot size) and scaled average gene expression (dot color) of cell type marker genes^93^ per cell type in the top panel, as well as the average gene score, calculated based on the chromatin accessibility of regulatory elements in the vicinity of the gene, of cell type markers in the bottom panel. **C)** Pseudo-bulk chromatin accessibility tracks scaled between 0-1 for endocrine marker genes by each endocrine cell type. **D)** Violin and boxplot depicting the distribution of the per-nucleus RNA (top) and ATAC (bottom) modality weights, divided by cell type. **E-F)** UMAP visualization of clustering based on the ATAC (E) or RNA (F) modality. **G)** Staged bar plot depicting the percentage (%) of cells in each cell type per condition.

The identified cell types were similarly represented in mice across conditions (**Figure 2G, Figure S2N, Table S9**) and correspond to the expected major cell types in islets in concordance with the literature^20^ with β-cells comprising the largest proportion of the endocrine cells (∼78-81%), followed by α-cells (∼7-9%), δ-cells (∼3-6%), and γ-cells (∼2-3%). In addition, we identified a small subset of endothelial, stellate cells and immune cells (∼3%, ∼0.3% and ∼1%, respectively), and only a low percentage of acinar cells (∼0.3-2%), indicating an overall high purity of the islet preparations with minimal exocrine contamination. The fraction of β-cells trends toward being increased with HFD feeding at the expense of α-cells and δ-cells.

### Short-Term High Fat Diet Induces a Proinflammatory Response in β-cells

To prioritize which cell types were most strongly affected by short-term HFD feeding, we used Augur^21^, a machine learning-based tool, which prioritizes cell types based on their molecular responses. Augur identified β-cells as the most strongly perturbed cell type (**Figure 3A**). Since β-cells are the most abundant cell type in the dataset, we evaluated whether the prioritization scores calculated by Augur were confounded by cell type proportions and found no significant correlation between the number of cells in each cell type and the Augur prioritization score **(Figure S3A)**. We also used differential expression analysis to prioritize cell types and identified the highest number of differentially expressed genes, totaling 6,101, between diet conditions in β-cells (**Figure 3B**). Although this method also prioritized β-cells as the most perturbed cell type, the result was strongly confounded by the number of cells in each cell type (**Figure S3B**). These results indicate that Augur is less biased than the number of differentially expressed genes for cell type prioritization in datasets with unbalanced cell type proportions, and that β-cells are most strongly affected by short-term HFD feeding.

**Figure 3:**
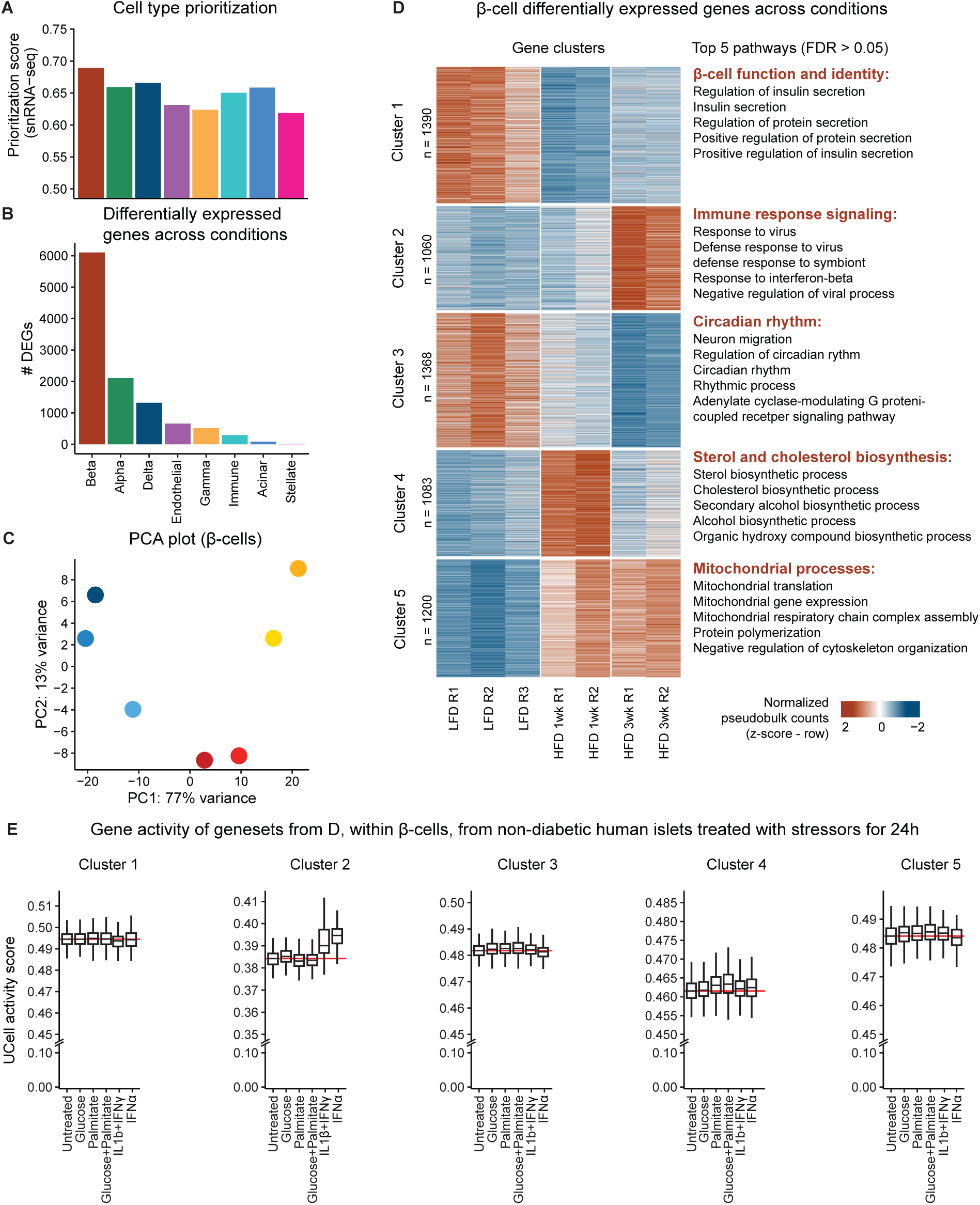
Transcriptional Dynamics of β-Cells in Response to One and Three Weeks of High-Fat Diet. **A)** Bar plot of Augur cell type prioritization scores across conditions, based on single-nucleus (sn)RNA-seq data, identified for each cell type. **B)** Bar plot of the number of differentially expressed genes identified for each cell type. **C)** Principal component analysis plot from the first two principal components calculated from pseudo-bulk snRNA-seq data from β-cells across all replicates, using the top 500 most variable genes. Each point represents a sample, with color and shading according to diet and replicate. **D)** Left panel: Heatmap showing z-transformed normalized pseudo-bulk average expression of differentially expressed genes identified within β-cells. Genes were divided into five clusters using fuzzy c-means clustering. Right panel: The top five most significant pathways from Gene Ontology biological processes within each fuzzy cluster. **E)** Boxplot of the per cell gene activity score of gene clusters from Figure 3D, within pancreatic β-cells from non-diabetic human donors which have been separated into groups treated with stressors; Glucose (22 mM) and/or palmitate (0.5 mM), interleukin-1beta (IL-1β) (1 ng/mL) and interferon gamma (IFNγ) (1000 U/mL), or interferon alpha (IFNα) (2000 U/mL) compared to untreated controls for 24 hours^26,27^. Red lines indicate the median in the untreated samples. Number of donors in each condition at each time point: 24hr; Glucose (n = 3), palmitate (n = 3), Glucose + Palmitate = (n = 3), IL-1β and IFNγ (n = 3), and IFNα (n = 2).

Principal component analysis of pseudo-bulked β-cell transcriptomes indicates high reproducibility between biological replicates, and a clear separation between diet conditions along the first principal component, explaining 77% of the variance in the dataset (**Figure 3C**). Using fuzzy clustering, we partitioned the 6,101 differentially expressed genes in β-cells into five clusters, with distinct temporal profiles, and performed pathway analysis (**Figure 3D**, **Figure S3C**, **Table S10-11**). Cluster 1 contains genes that are rapidly downregulated in response to HFD, and these are associated with β-cell identity (e.g. *Nkx6-1 and Mafa*), glucose sensing (e.g. *Slc2a2* and *Irs2*) and insulin secretion (e.g. *Glp1r*). In contrast, genes in cluster 2 exhibit a delayed upregulation and are associated with immune response signaling (e.g. *Stat1*, *Stat2*, *Irf7, Ifit1*, and *Cxcl10*). Specifically, the chemokine *Cxcl10* has been demonstrated to be secreted by islets from human organ donors with T2DM^22^. Mechanistic studies have shown that *Cxcl10* is enriched in pro-inflammatory extracellular vesicles secreted by β-cells. These vesicles contribute to induction of proinflammatory gene expression in neighboring β-cells and enhanced recruitment of CD8^+^ T-cells and macrophages^23^. Cluster 3 consists of genes that display sustained upregulation and is enriched for nuclear-encoded respiratory chain subunits of complex I (e.g. *Ndufs1* and *Ndufs3*), and genes involved in electron transport chain (ETC) complex IV assembly (e.g. *Smim20*, *Pet100*, and *Tmem223*). Genes in cluster 4 are transiently upregulated at one week of HFD and are associated with sterol and cholesterol biosynthesis (*Hmgcr*, *Sqle*, *Dhcr7*, *Cyp51*, and *Fdtf1*). Lastly, cluster 5 represents genes that are continuously downregulated, and include genes involved in transcriptional regulation of the circadian rhythm (e.g. *Clock*, *Per1/2*, *Cry1/2, and Rorc*). Circadian dysfunction has been implicated in the pathogenesis of T2DM, as islets from T2DM individuals show reduced expression of core clock genes such as *Per1* to *3* and *Cry1*^24^. Collectively, these analyses indicate that islet β-cells undergo rapid and diverse transcriptional changes in response to HFD, including a rapid decrease in the expression of genes related to β-cell identity, glucose sensing, and circadian rhythm, as well as a rapid induction of genes involved in mitochondrial function, sterol biosynthesis and a slower induction of many inflammatory genes.

Given the growing recognition of the role of inflammatory cytokines and nutrient stressors in β-cell dysfunction, and T2DM risk^25^, we investigated their involvement in the transcriptional response to compensation as induced by short-term HFD. To this end, we used publicly available scRNA-seq data of islets of Langerhans from non-diabetic human donors^26,27^. These islets had been treated *ex vivo* with various stressors, including diverse metabolic and inflammatory stressors in parallel for 24 or 72 hours. From each dataset, we extracted the β-cells and calculated gene module activity scores for each of the five clusters of DEGs (**Figure 3C and Figure S3C**). Glucose (22 mM), palmitate (0.5 mM), or glucose and palmitate together did not induce gene programs similar to any of the HFD-induced clusters in murine islets in this study, suggesting that glucose and palmitate, when added *ex vivo,* cannot recapitulate the effects of short-term HFD. In contrast, treatment with the pro-inflammatory cytokines interferon alpha (IFNα) (2000 U/mL), or the combination of interleukin 1beta (IL-1β) (1 ng/mL) and interferon gamma (IFNγ) (1000 U/mL) induced a gene program in human β-cells that is similar to cluster 2 in our dataset. Collectively, this indicates that short-term HFD induces a rapid pro-inflammatory response in β-cells.

### Pro-Inflammatory eRegulons are Activated in β-Cells Following Short-Term High Fat Diet

To identify transcriptional regulators of the rapid HFD-induced response in β-cells, we applied SCENIC+^28^, which infers gene-regulatory networks by identifying enhancer-driven regulons (eRegulons) containing transcription factors and their associated target enhancers and genes. We applied this approach to our paired snRNA-seq and snATAC-seq data and identified a total of 43 eRegulons. Filtering of these resulted in 16 high quality eRegulons, all of which were classified as activating eRegulons by SCENIC+ (**Figure 4A and Table S12-13**). High quality was determined by how well the expression of a transcription factor, and its motif activity, correlated with its activity scores from its potential target enhancers and genes across the different conditions (**Figure 4B, Figure S4, and Table S14**). Therefore, high-quality eRegulons are those where the transcription factors are active and expressed in conditions where their target genes are also expressed, and the target regions are accessible.

**Figure 4:**
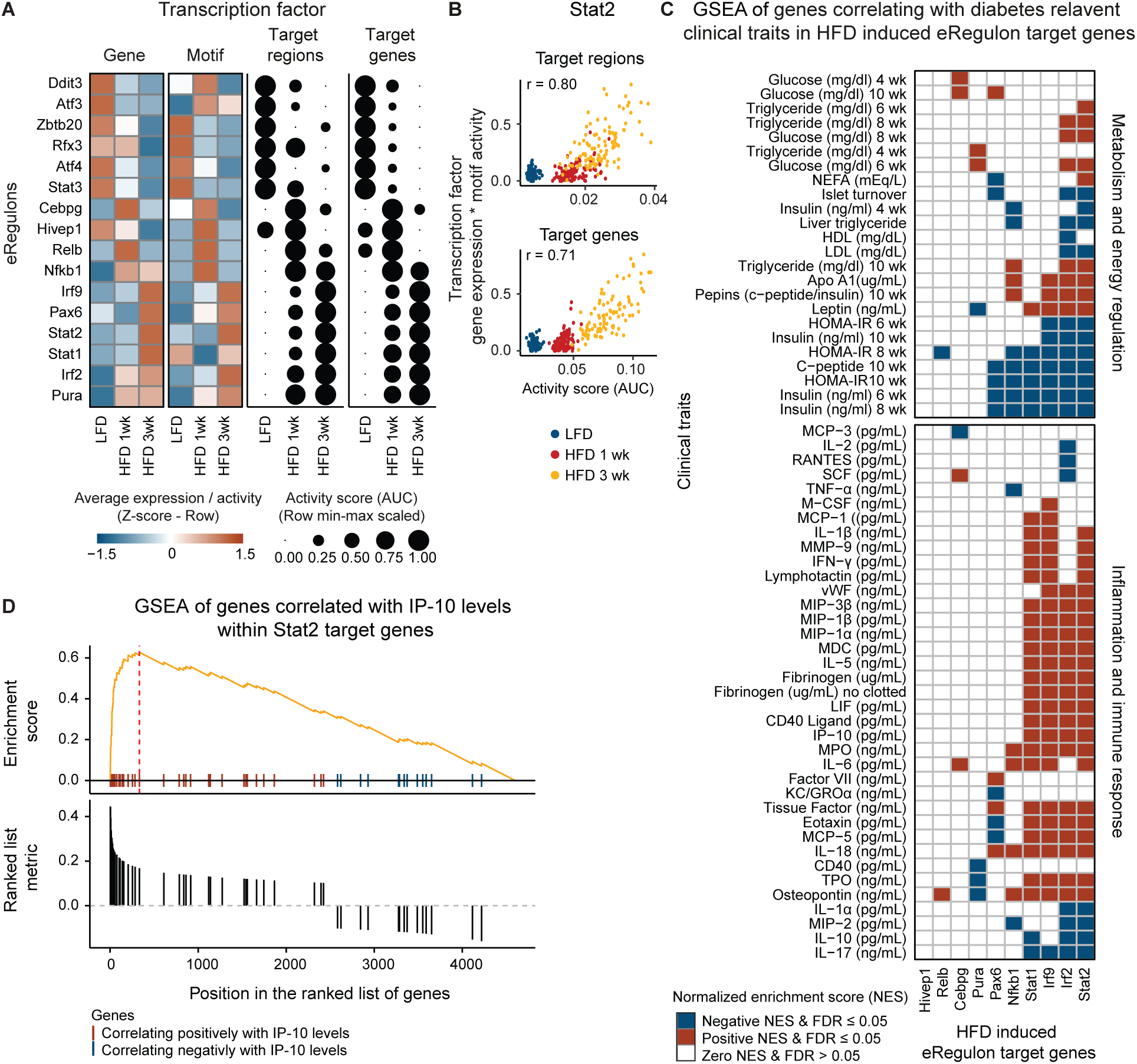
Immune and Inflammatory Transcription Factors Drive Transcriptional Changes in β-Cells in Response to Short-Term High-Fat Diet. **A)** Left panel: Heatmap displaying z-transformed normalized average gene expression (left) and motif activity (right) of eRegulon transcription factors across conditions. Right panel: Dot plot illustrating row min-max scaled average activity score (AUC) of target enhancers (left) and target genes (right), representing the gene set activity for each eRegulon per condition. Dot size corresponds to AUC. **B)** Pearson correlation analysis of semi-pseudo-bulked data (100 meta-cells per condition, each with 10 cells), from the Stat2 eRegulon. The y-axis depicts the min-max scaled product of semi-pseudo-bulked transcription factor, Stat2, gene expression, and motif activity. The x-axis depicts the activity score for Stat2 target enhancers (top) and Stat2 target genes (bottom). Points are colored according to condition (LFD: blue, HFD 1 week (wk): red, and HFD 3 wk: yellow). **C)** Heatmap illustrating gene set enrichment analysis (GSEA) of genes that correlate positively (correlation ≥ 0.1) or negatively (correlation ≤ -0.1) with clinical traits of diabetes from the Attie lab diabetes database (http://diabetes.wisc.edu/correl_f2.php) within high-fat-diet (HFD) induced eRegulon target genes. Genes associated with metabolism and energy regulation (top panel) as well as inflammation and immune response (bottom panel) are shown. Gene sets that are significantly enriched (false discovery rate (FDR) ≤ 0.05) within an eRegulon gene set are colored (blue if NES > 0 otherwise red). **D)** GSEA enrichment plot of the Interferon γ-induced protein 10 (IP-10, also known as CXCL10) gene set enriched within Stat2 target genes. Top: The y-axis represents the enrichment score, while the x-axis shows a rug plot of genes from the IP-10 gene set, found within the Stat2 target genes, ranked by their correlation with IP-10 levels, from high positive to low negative, and are colored based on their correlation score: positive (red) or negative (blue). The orange curve represents the running sum of the enrichment score across the ranked genes. The peak of this curve (indicated by the red dotted line) represents the maximum enrichment score for the gene set. Bottom: A bar plot illustrating the ranking of genes based on their correlation with IP-10 levels.

Following HFD feeding, several transcription factors, which have been implicated in β-cell function, display decreased gene expression and motif activity along with decreased target enhancer and target gene activity. These include Regulatory Factor X3 (*Rfx3*), which has been reported to play a role in islet development, and in the function of mature β-cells, including the regulation of glucokinase gene expression^29^.

In contrast to the downregulated eRegulons, the transcription factors driving upregulated eRegulons are not core regulators of β-cell identity and function. Instead, they are pro-inflammatory transcription factors, such as the NF-κB family (*Nfkb1* and *Relb*), which display a transient increase in expression, motif activity, and target enhancer and gene activity scores, as well as mediators of type I interferon (IFN) signaling, such as *Irf2*, *Irf9*, *Stat1,* and *Stat2*, which exhibit a delayed response to HFD, with some increase in target enhancer activity and target gene activity after one week of HFD. Type I IFN signaling is classically involved in innate antiviral responses but has also been implicated in the early stages of type 1 diabetes mellitus (T1DM) autoimmunity^30,31^, as well as in the dysregulation of insulin sensitivity and energy metabolism in diet-induced obesity in mice^32,33^.

We next explored whether the HFD-induced eRegulons are associated with diabetes-related clinical traits. To do this, we made use of the Attie lab diabetes database^34,35^, which includes bulk RNA-seq data of islets of Langerhans from obese B6:BTBR F2 mice, that either positively or negatively correlate with diabetes-related clinical traits. Genes with a Pearson correlation coefficient greater than 0.1 (positive correlation) or less than -0.1 (negative correlation) were included in the gene lists^34,35^. This means that a gene set that correlate positively with blood glucose levels indicates that increased expression of these genes is associated with higher blood glucose levels. Conversely, a negative correlation means that increased expression of these genes is associated with lower blood glucose levels or decreased expression of these genes is associated with high blood glucose levels. Gene set enrichment analysis (GSEA) revealed that the target genes of HFD-induced eRegulons are enriched in genes that either positively or negatively corelate with diabetes-related clinical traits. The analysis showed that the *Irf2*, *Irf9*, *Stat1,* and *Stat2* eRegulons were enriched for gene sets that correlate negatively with clinical traits related to insulin secretion, such as serum insulin and C-peptide levels, and *Irf2* and *Stat2* eRegulons are enriched for gene sets that correlate positively with plasma levels of glucose, non-esterified fatty acids (NEFA), and triglycerides. Furthermore, *Irf2*, *Irf9*, *Stat1,* and *Stat2* are also enriched for gene sets that correlate positively with circulating levels of chemokines, cytokines, and pro-inflammatory proteins, including for example interferon gamma-induced protein 10 (IP-10, *Cxcl10*), IL-1β, interleukin-6 (IL-6), Osteopontin, and IFNγ (**Figure 4C-D and Table S15**). These observations are concordant with the established role of these transcription factors in inflammatory signaling pathways^36–38^, and suggest a causal link between systemic pro-inflammatory molecules and regulation of β-cell function.

### β-Cells display a heterogenous response to inflammation

To identify potential sources of pro-inflammatory signals intrinsic to the islets, we used NicheNet^39^ to prioritize ligand-receptor pairs that most likely account for differential gene expression during LFD vs. one or three weeks HFD in β-cells, and to predict their downstream target genes. NicheNet identified tumor necrosis factor (TNF) and interferon beta (IFNβ) as immune cell-derived ligands within the top 30 prioritized interactions after both one and three weeks of HFD feeding (**Figure 5A and Table S16**). The 97 predicted target genes of TNF and IFNβ are all induced by HFD, and the majority reach maximum expression after 3 weeks of HFD feeding (**Figure 5B and Table S17**). Among the target genes are the pro-inflammatory transcription factors themselves, which suggests that there is a positive feedback loop that strengthens the inflammatory response.

**Figure 5:**
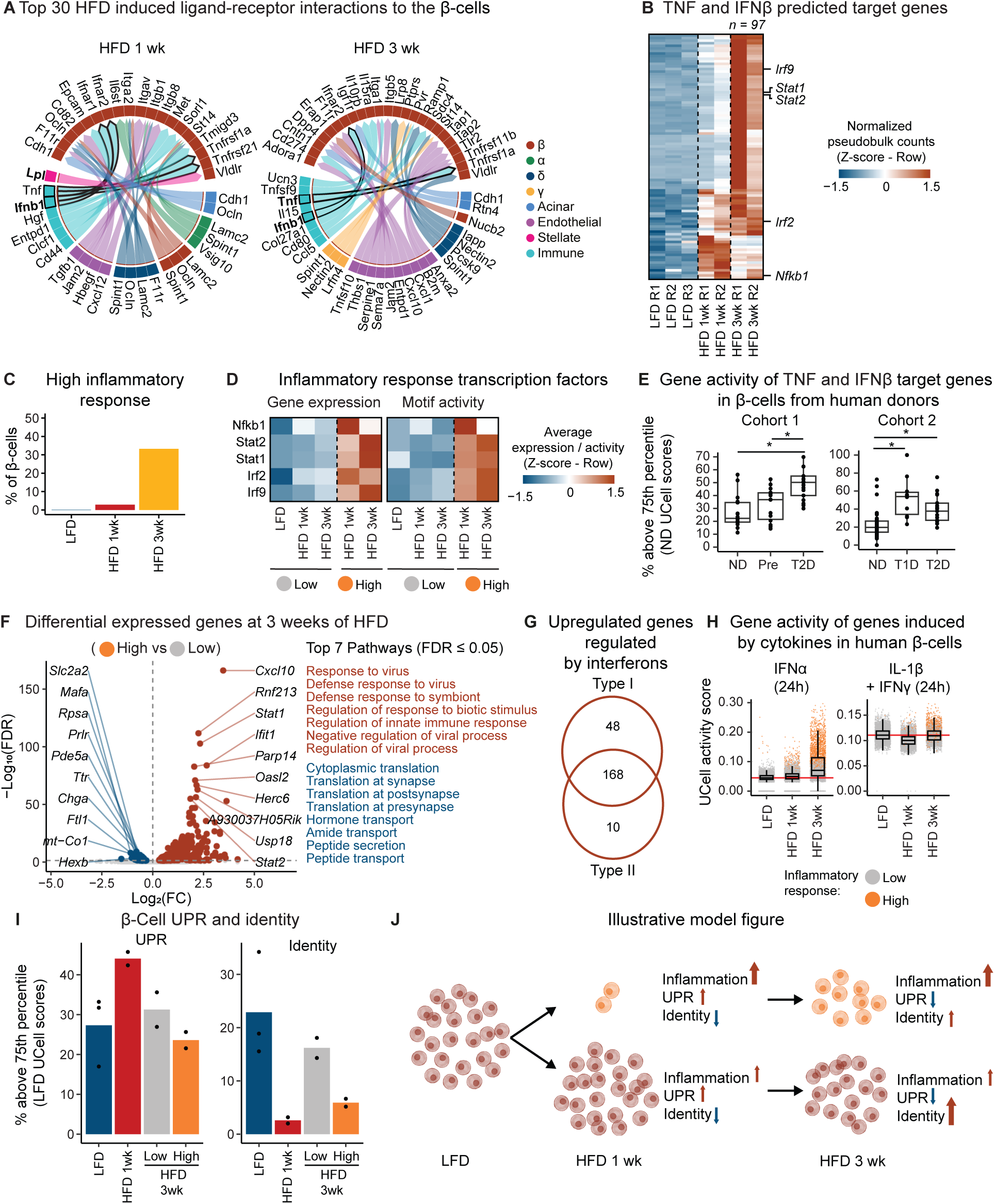
Identification of β-Cell subclusters based on Inflammatory Response. **A)** Circos plot of the top 30 predicted interaction links from all cell types to the β-cells from one week (1w) or three weeks (3wk) of high-fat diet (HFD). **B)** Heatmap displaying z-transformed normalized pseudo-bulk gene counts for putative tumor necrosis factor (Tnf) and interferon beta (ifnb1) target genes. **C)** Percentage of high inflammatory response β-cells at each condition. **D)** Heatmap displaying Z-transformed average gene expression (left) and motif activity (right) of inflammatory-related eRegulon transcription factors across conditions and β-cell low (grey dot) or high (orange dot) inflammatory subpopulation. **E)** Boxplot of the mean gene activity score for TNF and IFNβ target genes in β-cells from human donors, stratified by donor and disease status. Cohort 1^40^: Non-diabetics (ND, n = 17). Prediabetics (Pre, n = 14). Type 2 diabetics (T2D, n = 17). Cohort 2^40^: Non-diabetics (ND, n =38). Type 1 diabetics (T1D, n = 10). Type 2 diabetics (T2D, n = 17). Each point represents a donor. **F)** Left panel: Volcano plot depicting differentially expressed genes (DEGs) in high (orange dot) vs. low (grey dot) inflammatory response β-cell subpopulation at three-week HFD (HFD 3wk). The horizontal line indicates false discovery rate (FDR) = 0.05 and the vertical line indicates log2 fold change (FC) = 0. Top ten upregulated (red) and downregulated (blue) genes are annotated. Right panel: The top seven most significant (FDR ≤ 0.05) pathways from Gene Ontology biological processes for upregulated (red) and downregulated (blue) genes. **G)** Venn diagram showing results of enrichment search in INTERFEROME^42^ identifying an overlap between upregulated genes in high inflammatory β-cell subpopulation at three weeks of HFD, and genes known to be involved in interferon signaling pathways (type I, and II). **H)** Boxplot of the per β-cell gene activity score per condition using sets of genes that are significantly upregulated in pancreatic β-cells from non-diabetic human donors which have been treated with stressors; Interleukin-1beta (IL-1β) (1 ng/mL) and Interferon gamma (IFNγ) (1000 U/mL), Interferon alpha (IFNα) (2000 U/Ll) compared to untreated controls for 24h^26,27^. Grey dots represent β-cells in the low inflammatory subpopulation and orange dots represent β-cells in the high inflammatory subpopulation. Red lines indicate the median activity score in the LFD samples. **I)** Bar plot showing the mean prevalence of β-cells with high unfolded protein response (UPR) (left) or β-cell identity (right) as defined by the percentage (%) of cells above the 75th percentile of gene activity scores, using genes related to UPR or identity, within the low fat diet (LFD) β-cells, for each condition: LFD, HFD 1wk and HFD 3wk, where HFD 3 samples are subdivided by inflammatory state; low-inflammatory (Low) and high-inflammatory (High). Each point represents a replicate. **J)** Illustration of proposed mechanism: Following one week of HFD (HFD 1 wk) feeding UPR-mediated stress and transcriptional downregulation of β-cell identity genes in most β-cells, as well as induction of an inflammatory response in a small subset of β-cells (orange cells). After three weeks of HFD (HFD 3 wk), the UPR stress is resolved, but the subset of β-cells with an inflammatory response is increased. In β-cells without an inflammatory response, β-cell identity gene transcription is partially restored, but not in β-cells with an inflammatory response.

Using overrepresentation analysis, we compared activated predicted target genes of TNF and IFNβ ligand-receptor interactions with the target genes of HFD-induced eRegulons. We found the predicted TNF and IFNβ target genes to be significantly enriched (false discovery rate (FDR) ≤ 0.05) for genes predicted to be target genes of the inflammatory eRegulons driven by *Nfkb1*, *Irf2*, *Irf9*, *Stat1*, and *Stat2*, but not *Relb* (**Figure S5A and Table S18**). These results indicate that the transcriptional effects of TNF and interferon signaling are mediated by the activities of these transcription factors to modulate inflammatory gene expression patterns in response to short-term HFD.

There are conceptually two distinct mechanisms that can lead to an increase in the inflammatory response; either all cells have a modest response, or a subset of cells have a strong response. To explore which of these mechanisms are driving the inflammatory response in β-cells during short-term HFD feeding, we calculated a gene module activity score in every β-cell, using the TNF and IFNβ target genes. Generally, all β-cells in the HFD groups show slightly elevated scores, indicating that HFD influences the overall inflammatory response in β-cells across conditions. However, after three weeks of HFD, the distribution of these scores were right-skewed, suggesting that a subset of β-cells exhibit a pronounced inflammatory response at this time point (**Figure S5B**). By sorting the gene module activity scores for the inflammatory response gene set from all β-cells across HFD conditions, we determined the knee point of the curve as a threshold (gene module activity score ≥ 0.106) for segregating the β-cells into two subpopulations with high or low inflammatory response (**Figure S5C**). This analysis shows that there is an increase in the percentage of β-cells belonging to the high inflammatory subpopulation during exposure to HFD, from ∼0% under LFD conditions to ∼3% following one week, and ∼33% following three weeks of HFD (**Figure 5C**). Consistently, both the average gene expression and the inferred motif activity of the transcription factors *Nfkb1*, *Irf2*, *Irf9*, *Stat1*, and *Stat2* are higher in the high inflammatory sub-population compared to the low inflammatory subpopulation, which was not different from the LFD condition (**Figure 5D**).

To explore whether the inflammatory response observed in mouse islets in response to HFD is also observed in human β-cells, we utilized public scRNA-seq data from human non-diabetic subjects, prediabetes T2DM, or T2DM^40^ (Cohort 1), as well as data from individuals with or without T1DM or T2DM^41^ (Cohort 2). We utilized the TNF and IFNβ target genes to calculate the gene module activity score in every β-cell and used the 75^th^ percentile gene module activity score in non-diabetic β-cell as a threshold for defining cells with high inflammatory response (**Figure 5E and S5D**). In cohort 1, there was a tendency for an increased prevalence of high inflammatory β-cells in prediabetic compared to non-diabetic β-cells, though it was not statistically significant. In T2DM subjects, the increase was significant (p < 0.05). In cohort 2, the prevalence of high inflammatory β-cells was also significantly increased (p < 0.05) in both T2DM and T1DM β-cells, however to a much larger degree in the T1DM β-cells (**Figure 5E and S5D**). Collectively, this indicates that a subset of β-cells with a high inflammatory signature exists in humans and is related to the development of diabetes.

To explore the differences between high and low inflammatory response β-cells, we performed differential gene expression analysis between these two subpopulations at three weeks of HFD and found that 265 genes were significantly upregulated in the high inflammatory subpopulation, while 63 genes were downregulated (**Figure 5F and Table S19-20**). As expected, the genes upregulated in high relative to low inflammatory response β-cells are key regulators of interferon signaling, as well as genes related to antigen presentation, antiviral responses, immune cell recruitment, and immune checkpoint regulators (**Figure S5E**). Of the 265 upregulated genes, 226 were described in the INTERFEROME database^42^ as genes whose expression is induced by treatment with interferons (168 by both type I and II IFNs, 48 by type I, and 10 by type II) (**Figure 5G and Table S21-23**). To further support that this subset of β-cells is activated by type I IFN signaling, we used public data^26,27^ to define genes activated in human β-cells after acute type I IFN signaling (using IFNα) or after acute type II IFN signaling (using IL-1β and IFNγ). We observed a strong increase in gene module activity scores in high inflammatory response β-cells for type I IFN signaling but not type II IFN signaling (**Figure 5H and Figure S5F**).

The 63 downregulated genes, more highly expressed in low inflammatory β-cells, were associated with β-cell identity (e.g. *Mafa* and *Nkx6-1*), glucose sensing (e.g. *Slc2a2*), and insulin secretion (e.g. *Slc30a8* and *Chga*) (**Figure S5E**). Genes associated with β-cell identity and function was also expressed at higher levels in low inflammatory response β-cells at three weeks relative to one week of HFD (**Figure S5G**), thereby supporting previous studies indicating a transient reduction in the expression of these genes in islets after one week of HFD, followed by restoration at two and three weeks^14^. When insulin demand increases, β-cells may experience overwhelmed ER protein folding capacity, leading to misfolded proinsulin accumulation. To manage this stress, β-cells have been reported to activate an unfolded protein response (UPR)^6,43^. Comparison of gene expression in low inflammatory β-cells from LFD controls and one week of HFD revealed increased expression of the three key components of the UPR: *Atf6* (ATF6), *Ern1* (IRE1), and *Eif2ak3* (PERK)^44^ (**Figure S5H**). We identified a gene set for ATF6, IRE1, and PERK-mediated UPR (**Table S24**), and used the 75^th^ percentile gene module activity score in LFD β-cell as a threshold for defining cells with high UPR. The prevalence of β-cells with high UPR was transiently increased at one week of HFD, which returned to levels comparable to the LFD control after three weeks of HFD regardless of inflammatory state (**Figure 5I and Figure S5I-J**). Using the same approach to define β-cells with high expression of key β-cell identity genes (**Table S24**), we observed a transient reduction in the prevalence of β-cells with high identity after one week of HFD. However, as with the UPR response, this reduction was reversed by three weeks of HFD in low-inflammatory β-cells, with prevalence returning to LFD levels. In contrast, no such resolution was observed in high-inflammatory β-cells, where the prevalence was slightly elevated compared to one week of HFD.

Collectively, these results point to a mechanism where one week of HFD feeding increase insulin secretion from β-cells, which is accompanied by UPR-mediated stress and transcriptional downregulation of β-cell identity genes in most β-cells, as well as induction of an inflammatory response in a small subset of β-cells. After three weeks of HFD, the UPR stress is resolved, but the subset of β-cells with an inflammatory response is increased. In β-cells without an inflammatory response, β-cell identity gene transcription is partially re-stored, but not in β-cells with an inflammatory response (**Figure 5J**).

## Discussion

One of the hallmarks of the development of T2DM in humans is the transition from functional compensation to decompensation in pancreatic β-cells. In this study, we aim to identify the signals and regulators of this transition in a murine model of diet-induced obesity and insulin resistance by studying the effects of short-term HFD. At the physiological level, we observed that HFD had a rapid and significant impact on metabolic homeostasis consistent with previous research^14,15^. After one week of HFD, we observed hyperglycemia and impaired glucose tolerance relative to control mice as measured by IPGTT, indicative of insulin resistance. In addition, we observed a relative increase in both the secretory capacity of β-cells as measured by SPINA-Gβ and fasted serum insulin, indicating that β-cells partially compensate through increased insulin secretion. This functional compensation occurs in conjunction with strong induction of transcriptional programs related to UPR, cholesterol biosynthesis and oxidative phosphorylation, as well as repression of β-cell identity and function genes. After three weeks of HFD, we observed a reduction in the secretory capacity of β-cells relative to the controls as measured by SPINA-Gβ and a slight reduction in fasted serum insulin levels. These observations on the phenotypic level underscore that our model induces functional compensation, which transitions into early decompensation. To identify the signals and regulators of these functional transitions in islets of Langerhans, we mapped the transcriptome and the regulatory epigenome (measured through chromatin accessibility) using single nucleus multiomics.

Of all cell types in the islet of Langerhans, the β-cells undergo the largest transcriptional changes highlighting that β-cells play a pivotal role during functional transitions. Several recent studies have similarly reported decreased expression of genes associated with β-cell identity and function in both rodent and human β-cells in response to different metabolic stressors both *in vivo* and *in vitro*^45–49^. This has been suggested to be linked to the activation of UPR^50,51^, which may repress β-cell identity and function genes either through activation of transcriptional repressors, or through a mechanism known as squelching, where cofactors upon strong transcriptional activation of a novel gene set are redistributed from cell-type specific enhancers leading to ‘passive repression’^52^. Using our data, we were able to identify that activation of UPR (and other biological processes, including cholesterol biosynthesis, and oxidative phosphorylation) and repression of identity and function genes in β-cells occurs during the initial functional compensation phase *in vivo*. Thus, our observations support the hypothesis that β-cells secreting a large amount of insulin, simultaneously undergo transcriptional repression of identity and function genes mediated by UPR activation, oxidative and ER stresses through accumulation of misfolded proinsulin, and generation of reactive oxygen species due to high metabolic flow^14,43,44,53^.

After three weeks of HFD, we observed a decompensatory response as evidenced by a reduction in the secretory capacity of β-cells relative to the controls measured by SPINA-Gβ and a slight reduction in fasted serum insulin levels. Since the mice still display a relative reduction in glucose tolerance and hyperglycemia, indicative of insulin resistance, we believe this reduction in β-cell function represents β-cell exhaustion. This decompensation occurs in conjunction with the resolution of transcriptional induction of UPR.

Previous research suggest that the resolution of ER stress and UPR leads to normalization of transcription of β-cell identity and function genes^50^. In our data, we observed a dichotomous response, where approximately two-thirds of the β-cells restore the expression of identity genes to a large degree, whereas in the remaining β-cells, restoration of expression of identity genes is significantly blunted. The β-cells with blunted restoration of expression of identity genes upon UPR resolution are characterized by a strong proinflammatory response, which we map to IFN-α signaling.

Collectively, our observations point to a two-hit mechanism. In the early phase of insulin resistance, the increased demand for insulin is compensated by β-cells through increased insulin secretion, which simultaneously leads to transcriptional repression of β-cell identity and function genes potentially through squelching due to UPR activation and other gene programs. Upon β-cell exhaustion, the induced programs, including UPR, is resolved, but in a large subset of β-cells, a strong inflammatory response takes over and maintains the β-cell in a squelched state, thereby preventing normalization of expression of β-cell identity and function genes.

It has been established that obesity and T2DM in humans is associated with chronic lowgrade inflammation, characterized by increased circulating levels of cytokines and chemokines^54–56^, along with increased activation and recruitment of proinflammatory immune cells to multiple organs, including the endocrine pancreas^57–59^. Studies in mice have reported increased levels of circulating cytokines following just three days of HFD^60^, increased macrophage infiltration in murine islets after one week of HFD^61^ and in human islets in T2DM^58^.

In our data, we did not observe an increased number of immune cells in islets upon HFD, but caution is warranted due to the limitations of single-nucleus techniques in identifying rare cell types. For rare cell populations, the detection rate may not accurately reflect true proportions due to limited sampling or small variations during library preparation. However, based on inference of eRegulons, cell-cell communication analyses and association between metabolic traits and gene expression, we identified type I IFN signaling as the primary inflammatory signal in a subset of β-cells following short-term HFD feeding. Type I IFN signaling has been strongly associated with development of T1DM^30,62^, and is not well described in T2DM. We also found a fingerprint of these inflammatory processes being active and enriched in human β-cells in prediabetes and in both T1DM and T2DM, which suggests that activation of inflammatory responses in human β-cells may represent an important step in the transition from functional compensation to decompensation in early stages of insulin resistance. It remains unknown if the inflammatory signal we have identified in mice is conserved across species, or if other signals drive the inflammatory response in humans.

In conclusion, we have identified interferon-associated immune response as a key early event in HFD, driving inflammatory processes in pancreatic β-cells that may negatively impact cellular plasticity. These findings highlight the need for fine-grained, time-resolved investigations to dissect the early transcriptional mechanisms induced by HFD, as it remains unclear whether these inflammatory processes persist, intensify, or resolve beyond three weeks, and whether they ultimately exert detrimental or protective effects on cellular and metabolic health.

## Resource availability

### Load contact

Further information and requests for resources and reagents should be directed to and will be fulfilled by the lead contact Jesper Grud Skat Madsen.

### Materials availability

This study did not generate new unique reagents.

### Data and code availability

Raw and processed single-nucleus Multiome ATAC + gene expression sequencing data is available at NCBI GEO (https://www.ncbi.nlm.nih.gov/geo/) under accession number GSE293515. Furthermore, the supplementary file included in this paper provides detailed information on experimental diets (Table 1), phenotypic measurements (Tables 2–6), islet and nuclei recovery (Tables 7), cell type annotation (Tables 8–9), differential gene expression analysis (Table 10–11), eRegulons (Tables 12–15), cell-cell communications analysis (Tables 16–18), inflammatory subtype analysis (Tables 19–24), metadata (Tables 25–27), and reagents and tools used in this study (Table 29).

All code used used in this paper is available on Github: Phenotypic analysis (https://github.com/madsen-lab/islets_multiome_phenotyping), and Single-cell Multiome ATAC + Gene expression sequencing analysis (https://github.com/madsen-lab/islets_multiome_analysis).

Access for the public data used in this study has been described in the method section.

## Acknowledgements

Computation for this project was performed using the UCloud interactive HPC system, which is managed by the eScience Center at the University of Southern Denmark.

This manuscript used data acquired from the Human Pancreas Analysis Program (HPAP-RRID:SCR_016202) Database (https://hpap.pmacs.upenn.edu), a Human Islet Research Network (RRID:SCR_014393) consortium (UC4-DK-112217, U01-DK-123594, UC4-DK-112232, and U01-DK-123716). As well as data acquired from the Attie Lab Diabetes Database (RRID:SCR_016639) (http://diabetes.wisc.edu/).

This work was supported by grants from the Novo Nordisk Foundation (Advanced Grant), the Danish National Research Foundation (DNRF grant no. 141) to the Center for Functional Genomics and Tissue Plasticity (ATLAS).

## Author contributions

The project was conceptualized by I.V.S.E., S.M., and J.S.G.M. Animal work was carried out by I.V.S.E. and L.L., with assistance from A.M., S.M.N., and F.N. Bioinformatic analysis was conducted by I.V.S.E. under the supervision and guidance of J.G.S.M. Figures were prepared by I.V.S.E. with input from J.S.G.M. and S.M. The manuscript was written by I.V.S.E., J.G.S.M., and S.M. Funding for the project was acquired by S.M.

## Declaration of interests

The authors declare no competing interests.

## Method Details

### Mice

8-week-old C57BL/6J male mice from Taconic Bioscience (C57BL/6JBomTac; B6JBOM-M) were kept in IVC-cages, up to 4 mice per cage, and maintained in a 12-h light–dark cycle, at room temperature of 22 ± 3 °C and ∼55% humidity with ad libitum access to food and water. The mice were acclimatized for three weeks and fed LFD (Research Diets; D12450J) prior to experiment start. Hereafter, at day 0, the 11-week-old mice were divided into three groups. One group continued the LFD as a control, and one group were fed either HFD (Research Diets; D12492) for one week or three weeks. At the end of feeding, the mice were used for several studies, including phenotypically characterization and Single-cell Multiome ATAC + Gene expression sequencing as described below.

### Intraperitoneal Glucose Tolerance Test

After one week or three weeks, the glucose tolerance of the mice was assessed by intraperitoneal glucose tolerance test (IPGTT). Mice were fasted for 4 hours in the morning and fasting plasma blood glucose levels were determined (T = 0). D-(+)-Glucose (Sigma-Aldrich; G8270) was administered intraperitoneally (2 g/kg) in a 0.9% saline suspension (B. Braun; 420079) and plasma blood glucose levels were measured at 15, 30, 60 and 120 min after glucose injection using a glucometer (FreeStyle Freedom Lite; Cat#70919-70).

### Intraperitoneal Insulin Tolerance Test

After one week or three weeks, the insulin tolerance of the mice was assessed by intraperitoneal insulin tolerance test (IPITT). Mice were fasted for 4 hours in the morning and fasting plasma blood glucose levels were determined (T = 0). Insulin (Actrapid; A10AB01) in 0.9 % saline suspension (B.Braun; 420079) was administered intraperitoneally (0.75 U/kg) and plasma blood glucose levels were measured at 15, 30, 60 and 120 min after insulin bolus using a glucometer (FreeStyle Freedom Lite; 70919-70).

### Fasting Serum Insulin

After one week or three weeks, the mice were fasted for 4 hours in the morning. Tail or hindleg blood samples were collected and allowed to clot at room temperature for 30 min, whereafter the samples were centrifuged for 20 min at 2,000 x g at 4 °C. The serum supernatant was extracted and used for insulin quantification using the Ultrasensitive Mouse Insulin ELISA kit (Chrystal Chem; 90080) according to the manufacturer’s instructions.

### Estimating Insulin Sensitivity and β-cell Function

We used a mathematical model of insulin-glucose homeostasis called SPINA (Structure Parameter Inference Approach)-Carb^63^ to estimate two key parameters: insulin receptor gain (SPINA-GR), which measures insulin sensitivity, and β-cell insulin secretory capacity (SPINA-Gβ), which measures β-cell function. Both parameters were calculated using an implementation of the SPINA-Carb model (Version 5.0.0)^64^ in an R statistical environment. The calculations relied on steady-state fasting concentrations of serum insulin (pmol/L) and blood glucose (mmol/L), employing the functions SPINA.GR for insulin receptor gain and SPINA.GBeta for β-cell secretory capacity.

### Statistical Analysis

Data from phenotyping experiments: IPIGTT, IPITT, fasted blood plasma glucose, fasted serum insulin, body weight gain, SPINA-GR, and SPINA-Gβ were analyzed using the non-parametric Wilcoxon test to compare two means.

All code and parameters used for this analysis are available on GitHub (https://github.com/madsen-lab/islets_multiome_phenotyping). All analysis was carried out in Rstudio (version 4.4.0).

### Islet Isolation

Mice used for islet isolation were sacrificed using cervical dislocation. The pancreas was injected through the common bile duct with 3 mL dissociation buffer (0.7 mg/mL Collagenase P (Merck; Lot#33768602), 1X HBSS (TermoFisher; 14065049) and 0.3 g/L NaHCO3 (Sigma-Aldrich; S5761)), using a 30G syringe. The tissue was dissociated in a tube with 2 mL dissociation buffer for 15 min in a 37 °C water bath. To further dissociate the tissue, the tubes were shaken gently by hand for 1 min. The dissociation was quenched by adding cold quenching buffer (1X HBSS, 0.3 g/L NaHCO3 and 10 % heat-inactivated fetal bovine serum (VIA) (Biochrom; B15-005)) to a total volume of 20 mL.

The tissue solution was centrifuged for 2 min at 150 x g at room temperature (RT) in a swing-bucket centrifuge (acceleration = 8, brake = 2), the supernatant was aspirated until there was 10 mL left. This wash step was repeated two times. The tissue was further dispersed by pipetting the tissue solution through a 14G venflon (without the needle) 5 times. Hereafter, the tissue solution was spun down for 7 min at 200 x g at RT, supernatant removed, and tissue pellet was resuspended in 10 mL of a histopaque density gradient used to separate islets from non-endocrine tissue (5:11 Histopaque-1077 (Sigma-Aldrich; 10771) and 6:11 Histopaque-1119 (Sigma-Aldrich; 11191)). The Histopaque solution was centrifuged for 20 min at 380 x g at RT (Acceleration = 8, break = 2) in 15 mL tubes. The islet containing supernatant was aspirated and the supernatant was filtered through a 400 μm filter (pluriStrainer; 43-50400-03), followed by reversed filtering using a 70 μm filter (Fisherbrand; 11597522). Islets were hand-picked two times and contained in RPMI 1640+ GlutamaxTM medium (Gibco; 61870044) (11 mM glucose) supplemented with 5 % heat inactivated fetal calf serum (Biochrom; B15-005) and 100 U/mL penicillin and 100 mg/mL streptomycin (Lonza; DE17-602E).

### Nuclei Isolation from Islets

We adapted the 10x Genomics protocol (CG000375, Demonstrated Protocol_NucleiIsolationComplexSample_ATAC_GEX_Sequencing_RevB)^65^ for isolation of nuclei from isolated islets. Detergent-based permeabilization of the cell membrane was used for nuclei isolation. Freshly isolated islets were moved to biomasher II Eppendorf tubes and centrifuged for 5 min at 400 x g in a swing-bucket centrifuge at 4°C. The media-supernatant was carefully removed, leaving ∼ 20 µL media on top of the islet pellet. To release the nuclei from the islets, 100 μL nuclei preparation buffer (NBP) (10 mM HEPES pH 7.5 (Sigma; H3375), 1.5 mM MgCl2 (Sigma;M8266), 40 mM KCl (Sigma; P9541), 250 mM sucrose (Sigma; S0389), 0.5 mM Spermidine (Sigma, S2626), 1% BSA (Merck; M0314L), 0.1% IGEPAL CA-360 (Sigma;I8896), 0.2 mM DTT (NEB, B1034A) and 1 U/μL RNase inhibitor (NEB; M0314) dissolved in DEPC water) was added to the islet pellet and the pellet was homogenized 20 x times using the Biomasher II Disposable Micro Tissue Homogenizer (EOG-sterilized) (NIP; 320103). Hereafter, 1000 μL NBP was added to the tube. The solution was incubated on ice for 5 min, gently pipetting halfway using wide-bore pipette tips. To remove cell debris, the nuclei suspension was passed through a 70 μm tip strainer into a 1.5 mL DNA low bind Eppendorf tube. To increase the yield of nuclei the biomasher Eppendorf tube was washed with 500 μL NPB, and passed through the tip strainer, into the same 1.5 mL tube. The nuclei suspension was centrifuged at 500 x g in a swing-bucket centrifuge for 5 min at 4 °C and the NPB supernatant was removed until there was ∼50 μL left. The nuclei suspension was washed by adding 1000 μL 2 % BSA in 1X PBS + 1U/μL RNase inhibitor to the nuclei suspension (not mixed!) and after 5 min on ice, the suspension was mixed to resuspend the nuclei pellet. Hereafter, the nuclei suspension was centrifuged at 500 x g in a swing-bucket centrifuge for 5 min at 4 °C, and the BSA supernatant was removed.

To permeabilize the nuclei membrane for transposition the nuclei were incubated in 100 μL 0.1X lysis buffer (10 mM Tris-HCl pH 7.5 (VWR; 103156X), 3 mM MgCl2, 10 mM NaCl (Sigma; S3014), 0.01 % Tween-20 (Sigma; P1379), 0.01 % IGEPAL CA-630, 1 % BSA, 0.001 % Digitonin (ThermoFisher; BN2006), 1 mM DTT, 1U/μL RNase inhibitor dissolved in DEPC water). After 2 min the permeabilization was stopped by diluting the lysis buffer with 1 mL Wash buffer (10 mM Tris-HCl pH 7.5, 3 mM MgCl2, 10 mM NaCl, 0.1 % Tween-20, 0, 1 % BSA, 1 mM DTT, 1U/μL RNase inhibitor dissolved in DEPC water). The solution was mixed by careful pipetting 5 times. Samples were centrifuged at 500 x g in a swing-bucket centrifuged for 5 min at 4 °C and the wash buffer supernatant was removed. The nuclei pellet was resuspended in 60 μL 1X nuclei buffer (10x Genomics), 1 mM DTT and 1 U/μL RNase inhibitor dissolved in DEPC water). Finally, any nuclei aggregates were removed by passing the nuclei solution through a 40 μm tip strainer into a new DNA low bind Eppendorf tube, and the nuclei suspension was centrifuged at 500 x g in a swing-bucket centrifuge for 5 min at 4 °C and the supernatant was removed until there was 7 μL nuclei buffer left.

For counting, 1 µL nuclei solution was diluted in 10 µL 1X nuclei buffer and 9 µL trypan blue dye (Bio-rad; Cat#1450021). Nuclei were counted using a Bürker pattern counting chamber (Sigma-Aldrich, BR718920). 5000-16,000 nuclei were submitted for Single-cell Multiome ATAC + Gene expression sequencing.

### Single Cell Multiome ATAC + Gene expression sequencing

We used the Single-cell Multiome ATAC + Gene expression sequencing technique provided by 10X Genomics, to simultaneously detect changes in gene expression (snRNA-seq) and chromatin accessibility (snATAC-seq) .This approach was applied to nuclei from islets of Langerhans isolated from seven mice (143-300 islets / mouse) fed with either LFD (n = 3) or HFD for either one week (n = 2) or three weeks (n = 2). Each mouse sample was processed separately.

Isolated nuclei were prepared for transposition and library generation using the following 10x Genomics kits: Chromium Next GEM Single Cell Multiome GEM Kit A, 16 rxns (PN-1000232), Chromium Next GEM Single Cell Multiome Amp Kit A, 16 rxns (PN-1000233), Chromium Next GEM Single Cell Multiome ATAC Kit A, 16 rxns (PN-1000280), and Library Construction Kit, 16 rxns (PN-1000190).

The samples were prepared in accordance with the manufacturer’s instructions (CG000338, Chromium Next GEM Single Cell Multiome ATAC + Gene Expression User guide). Libraries were sequenced on the Illumina NovaSeq 6000 System (Illumina, 20012850).

### Single Cell Multiome ATAC + Gene expression sequencing Analysis

#### Demultiplexing

Raw base call files were demultiplexed into FASTQ files using the mkfastq pipeline in 10X Genomics Cell Ranger ARC (version: 2.0.1)

#### Generation of count matrices

All code and parameters used for this analysis are available on GitHub (https://github.com/madsen-lab/islets_multiome_analysis). All analysis except for alignment and SCENIC+ analysis was carried out in Rstudio (version 4.3.0).

snRNA-seq data was aligned to the mm10 (GRCm38) reference genome with STARsolo (version: 2.7.9a)^66^, counting both intronic and exonic reads or only exonic reads (soloUMIlen = 12, soloCellFilter = none, soloType = CB_UMI_Simple, soloCBmatchWLtype = 1MM, clipAdapterType = CellRanger4, soloFeatures = GeneFull_Ex50pAS Gene). The snATAC-seq data was aligned to the mm10 reference genome using cellranger-arc count (default parameters) from 10X Genomics Cell Ranger ARC (version: 2.0.1)^67^.

#### Quality control

The snRNA-seq count matrix was filtered for high-quality barcodes using valiDrops (version: 0.1.0)^68^ functions rank_barcodes, quality_metrics, and quality_filter (default parameters). Additionally, we added a global filter for the mitochondrial fraction < 0.025 and exon ratio > 1. The snATAC-seq count matrix was filtered using ArchR (version: 1.0.2)^69^ (TSSEnrichment ≥15, BlacklistRatio ≤ 0.05, nFrags ≥ 2500). Doublets were identified in both modalities using DoubletFinder (version: 2.0.3)^70^ for the RNA modality and ArchR for the ATAC modality. Nuclei which passed quality control in both modalities were retained. Nuclei called as doublets in both modalities were removed.

#### Dimensional reduction and cluster identification

Weighted nearest neighbor analysis (WNN) (version: 4.4.0)^19^ was used to generate a single representation of the combined snRNA and snATAC-seq data. WNN analysis requires two dimensional reductions, one for each modality.

Dimensional reduction for the snRNA-seq data was computed in seurat workspace. Count normalization, variable feature selection, dimensional reduction and batch integration was performed simultaneously using JOINTLY (version: 1.0) (default parameters)^71^. snRNA-seq counts were normalized with Seurat’s LogNormalize function (scale.factor=1e4)

Before processing the snATAC-seq data, the snRNA-seq data was used to further identify any remaining doublets based on poly-hormone expression of the major endocrine marker genes: *Ins2*, *Gcg*, *Sst*, and *Ppy*. Using a deep clustering approach (resolution = 20), clusters were identified as potential doublets (polyhormonal clusters) if they had an average scaled expression above 0.6 for more than one of marker gene with a sum of scaled expression values exceeding 1. These were removed from both modalities.

Hereafter, a dual-fold methodology was employed to compute batch-corrected dimensional reductions for the snATAC-seq data. Variable feature selection and integration was performed using 5000 bp window tiles. The top 150,000 most variable windows were identified using ArchR’s Iterative latent semantic indexing (LSI), with 4 iterations, these were used for integration with LIGER (version: 1.0.0)^72^ (default parameters). The resulting embeddings were used to cluster the snATAC-seq data (20 dimensions, resolution = 2), for putative cluster annotation which was used for peak detection. The clusters were annotated by evaluating the gene score (a prediction of gene expression based on the accessibility of regulatory elements in the vicinity of the gene) and gene expression of major marker genes. We used the imputed weight method MAGIC on the gene scores for reducing noise of the snATAC-seq data. The identified clusters were subsequently used for reproducible peak calling using MACS2 in ArchR^73–75^ (minCells = 20, maxCells = 5205, minReplicates = 2, maxReplicates = 3). Lastly, the generation of batch-corrected dimensional reductions using LIGER was repeated as described above, but now using the top 150,000 variable peaks instead of windows.

Meta data information, peak matrix, gene scores and LIGER embeddings from the ArchR object were transferred to the seurat object containing the snRNA-seq data using ArchRtoSignac (version: 1.0.3)^76^. From here, analysis was performed using the combined data in the seurat object unless otherwise specified.

A WNN-graph was constructed using both modalities (rna embeddings = 1:15, peak embeddings = 1:20) and used for graph-based clustering as well as Uniform Manifold Approximation and Projection (UMAP) visualization.

Cell type identities were manually assigned to cell clusters by evaluating expression of prior knowledge cell-type markers.

#### Evaluation of batch mixing

Batch mixing of replicates before and after integration was evaluated for each diet condition in both modalities. This evaluation was conducted using unintegrated snRNA-seq principal component (PC) embeddings (1:15 dimensions) and snATAC-seq singular value decomposition (SVD) embeddings (1:20 dimensions), as well as integrated snRNA-seq JOINTLY embeddings (1:15 dimensions) and snATAC-seq LIGER embeddings (1:20 dimensions).

The assessment employed the local inverse Simpson’s index (LISI)^77^ and the average silhouette width (ASW). Integration LISI (iLISI) and batch ASW (bASW) were calculated using these embeddings with the evaluateEmbedding function from (http://www.github.com/madsen-lab/JOINTLY_reproducibility)^71^.

Both metrics were rescaled as previously described by Luecken, M.D. et al.^78^. For iLISI, a value of 0 indicates low batch integration, while a value of 1 indicates high batch integration. For bASW, a value of 0 reflects low cell type separation and high batch separation, whereas a value of 1 indicates optimal performance with high cell type separation and low batch separation.

### Cell type prioritization

To identify the cell type in which transcription was the most perturbed by HFD feeding, we performed cell type prioritization analysis using Augur (version: 1.0.3)^21^ using the calculate_auc function (subsample_size = 12, n_subsamples = 150). The most perturbed cell type is the cell type with the highest area under the curve (AUC) exceeding 0.5.

### Differential gene expression analysis

Differential gene expression analysis was performed using the DESeq2 package (version: 1.40.2)^79^ between condition and clusters (including subclusters), using aggregated (pseudo bulk) counts from each cell type per replicate obtained with Edgers Seurat2PB function (version: 3.42.4)^80^ (default parameters). The DESeq function (default parameters) was utilized to estimate size factors, count normalization and calculating differential gene expression based on a negative binomial distribution. Comparisons across diet was performed using the Likelihood Ratio Test (LRT) to identify genes, which change across time points i.e. genes with a dynamic expression pattern. Pairwise comparisons between overall cell type clusters as well as between subclusters were performed using a paired Wald test, specifying which condition each replicate is from. Genes were filtered based on their adjusted p-value (FDR ≤ 0.05).

### Fuzzy clustering of genes

To explore the temporal regulation of gene expression, all genes with a dynamic expression pattern (i.e. those changed in any time point across conditions, identified in the DESeq2 LRT analysis), were clustered based on normalized pseudo bulk gene expression averaged by condition, using fuzzy clustering with the package MFuzz (version: 2.60.0)^81^. First, expression values were standardized using the standadise function (default parameters), and the optimal fuzzifier was estimated using mestimate (default parameters). Hereafter, the optimal number of clusters for the identified fuzzifier was performed by calculating the minimum distance between 1 to 20 clusters by using the function Dmin (crange=seq(2,20,1)), repeats = 15). The optimal number of clusters was defined as the number of clusters at which the minimum centroid distance begins to decrease slowly. Finally, clustering was performed with mfuzz (default parameters). The genes in each cluster were used for pathway analysis.

### Pathway Analysis

Pathway analysis was performed to explore the functional relevance of differential expressed genes. The analysis was performed using the function enrichGO (ont = “BP”, pvalueCutoff = 0.2, qvalueCutoff = 0.2) from the ClusterProfiler package (version: 4.8.3)^82^. Mouse genes were annotated using the org.Mm.eg.db package (version: 3.17.0)^83^. GOterms for biological processes were used as reference gene sets, and all genes which were used as input for the DESeq2 analysis were used as background (universe). Gene sets with an FDR ≤ 0.05 were considered significantly enriched.

### eRegulon identification

In Python (version: 3.8.17), eRegulon interference analysis in β-cells across conditions was performed with SCENIC+ (version: 1.0.1.dev4+ge4bdd9f)^28^. Raw snRNA-seq counts, peak matrix, and condition annotations (for nuclei within the β-cell cluster) were extracted from the seurat object and utilized for the analysis. First, PyCisTopic (version: 1.0.3.dev18+ge563fb6)^84^, within the SCENIC+ framework, was used to perform latent Dirichlet allocation topic modeling on the peak matrix (n_itr = 500, alpha = 50) to identify sets of co-accessible regions (topics). Subsequently, PyCisTarget (version: 1.0.3.dev1+g3fde1ce)^28^, within the SCENIC+ framework, was utilized to identify differentially accessible regions (DARs) using a Wilcoxon rank-sum test (default parameters) and to identify enriched motifs within the topics and DARs (default parameters). Motif enrichment analysis was carried out using the clustered motif collection database from SCENIC+ (v10, mm10). The results, together with the raw snRNA-counts, were used as input to the SCENIC+ algorithm (default parameters) against a list of all known mouse transcription factors provided by SCENIC+. Identified putative eRegulons were then employed for downstream analysis.

### Identification of High Quality eRegulons

High-quality eRegulons were identified through correlation analysis of semi-pseudo-bulked data per condition (100 meta-cells per condition, each with 10 cells) using Pearson correlation. The correlation analysis was performed using the min-max scaled product of transcription factor gene expression and of transcription factor motif activity with activity scores for target genes and enhancers from SCENIC+ (computed using AUCell^85^), for each eRegulon. This was to ensure that transcription factors were active in cells where their target genes were expressed, and target regions were accessible (correlation >= 0.6).

Motif activity analysis, used in the correlation analysis, was performed using chromVar (version: 1.22.1)^86^ (default parameters) within the ArchR framework, using the motif collection database from SCENIC+. Subsequently, the motif activity for each of transcription factor was calculated by averaging the deviation score, from chromVar, for each motif associated with the specific of transcription factor.

### Gene Set Enrichment Analysis (GSEA)

GSEA was performed with a focus on determining whether eRegulon target genes were enriched within genes associated with clinical phenotypes of diabetes. These genes were derived from the Attie lab diabetes database (http://diabetes.wisc.edu/correl_f2.php, access: November 15^th^, 2023)^34,35^, who has performed correlation analysis between diabetes-related clinical phenotypes and gene expression within islets of Langerhans, from obese B6:BTBR F2 mice. For each clinical phenotype, all genes with a Pearson correlation coefficient > 0.1 (positive correlation) or < -0.1 (negative correlation), were used to create gene lists. The gene lists for each clinical phenotype were ranked in descending order of their correlation coefficients. This ranked list was then used as input for the GSEA, while the eRegulon target genes acted as the pathways being tested for enrichment. GSEA was performed using clusterProfiler (version: 4.8.3)^82^ (default parameters). The analysis was performed for each clinical phenotype, specifically focusing on phenotypes categorized under “Metabolism and Energy Regulation” and “Inflammation and Immune Response.” Only phenotypes that yielded at least one significant enrichment hit (FDR ≤ 0.05) were retained for visualization.

### Cell-Cell communication analysis

To infer cell-cell communication between β-cells and all other cell types across conditions (LFD, one- and three-week HFD), we used the MultiNicheNet package (version: 1.0.3) ^39^, to predict ligand-receptor interactions between all cell types (senders) and β-cells (receivers) between conditions. First, MultiNicheNet uses aggregated pseudo bulk counts and the DESeq2 package for differential gene expression between conditions. Differential expressed ligands, receptors, and target genes were identified using the thresholds: logFC > 1 and adjusted p-value ≤ 0.05. The ligand activity within receivers was calculated using the top 250 target genes with the highest regulatory score (this score reflects how much prior knowledge supports specific ligand – target gene regulation). Ligand-receptor interactions were prioritized, within each condition, using a weighted combination of min-max scaled scores of the following criteria as recommended by the developers: 1. upregulation of the ligand in the senders and upregulation of the receptor in the receiver at the condition of interest (weight = 1), 2. sufficient expression levels of the ligand and receptor in samples of the same group (weight = 1, to mitigate influence of outliers), 3. cell type and condition specific expression of the ligand in the sender, and receptor in the receiver (weight = 2, to mitigate the influence of upregulated ligand / receptors that are weakly expressed), and lastly, 4. high ligand activity (weight = 2). Based on this combined score, the top 30 ligand-receptor interactions, for each condition, were visualized in a circos plot. Target genes of ligand-receptor pairs were identified by calculating Pearson and Spearman correlation coefficients between the min-max scaled product of pseudo bulk expression of the ligand and receptor of interest and the expression of the target gene. Target genes with a correlation coefficient greater than 0.5 or less than -0.5 were used for downstream analysis.

### Overrepresentation analysis of gene sets (eRegulon target genes and ligand target genes)

Overrepresentation analysis of gene sets was performed by constructing a 2 × 2 contingency table. This analysis aimed to explore the potential association between gene sets, denoted *S* (such as target genes identified in cell-cell communication analysis) and eRegulon target genes, denoted D, and *N* is the total number of genes:

| | In $S$ | Not in $S$ |
| --- | --- | --- |
| In $D$ | $N_{DS}$ | $N_D - N_{DS}$ |
| Not in $D$ | $N_S - N_{DS}$ | $N - (N_D + N_S - N_{DS})$ |

Here *N_DS_* is the total number of genes in both *D* and *S*, *N_D_-N_DS_* is the number in genes that are in *D* but not in *S*, *N_S_-N_DS_* is the number of genes that are in *S* but not in *D*, and *N*-(*N_D_+N_S_-N_DS_*) is the number of genes that are in neither *D* nor in *S*. Fisher’s exact test was employed to determine if there was a significant association between *D* and *S*. Significant associations was determined based on adjust p-values (FDR ≤ 0.05).

### Gene Module Activity

A ranked based methods UCell (version: 2.4.0)^87^ (maxRank = 1000) were used to quantify the activity of gene sets that were not calculated in SCENIC+ (i.e., eRegulon target genes and enhancers). UCell calculates activity scores by ranking all genes (up to the maximum rank) expressed in each individual cell based on their expression levels, where a lower rank indicates greater expression. For a gene set of interest, UCell performs a Mann–Whitney *U* test, with the activity score represented as the normalized *U*-statistic, which ranges from 0 to 1. A score of 1 indicates that the genes in the gene set are among the most highly expressed in the cell, while a score of 0 indicates that these genes are among the least expressed.

### Inflammatory Response Subcluster identification

For the identification of β-cells with a high inflammation, the TNF and IFN-β target genes identified by MultiNicheNet, were utilized to define an “ Inflammatory response” gene set. Subsequently, UCell activity scores were computed for each β-cell, following the procedure outlined in the section “Gene Module Activity”. Activity scores for β-cells from the HFD week one or HFD week three conditions were sorted in a descending order, and used to generate a knee point curve, with activity scores on the y-axis and number of β-cells with that activity score on the x-axis. The inflection point of this curve, indicative of a significant change in the trend, was determined using the inflection package (version: 1.3.6)^88^ (default parameters).

β-cells with an activity score exceeding the inflection point (>0.106) were annotated as belonging to a high inflammatory subcluster.

### Public Single-cell RNA-seq Data

To identify genes induced by glucose, palmitate, or cytokine treatment in human β-cells, we sourced scRNA-seq data from human non-diabetic pancreatic islet cells treated with glucose (22 mM) and/or palmitate (0.5 mM), IL-1β (1 ng/mL) and IFNγ (1000 U/mL), IFNα (2000 U/mL), or untreated as control for 24 hours and 72 hours^26,27^. Filtered barcode-, gene - count – matrices, and a meta data file was available for download from GEO (GSE218316) and used to create a seurat object. The seurat object was subset to only include β-cells based on the provided annotation “Celltype”. Differentially expressed genes were identified between treated and untreated β-cells following 24 or 72 hours of treatment using pseudo-bulked counts, and a DESeq2 paired Wald test as described in the section “Differential gene expression analysis”. We filtered genes with an adjusted p-value (FDR ≤ 0.05) and a log2 fold change (log2FC ≥ 0.5) to retain those with significantly increased expression after treatment. Subsequently, these genes were converted from human gene symbols to mouse gene symbols, by mapping human gene symbols to Entrez IDs using AnnotationDbi (version: 1.62.2)^89^ and org.Hs.eg.db (version: 3.17.0)^90^. We then matched these human Entrez IDs to mouse orthologs using Orthology.eg.db (version: 3.17.0)^91^ and converted the mouse Entrez IDs to mouse gene symbols using org.Mm.eg.db (version: 3.17.0)^83^. Subsequently, these genes were then used to compute gene module activity scores for each β-cell, following the procedure outlined in the section “Gene Module Activity”.

Furthermore, to assess whether inflammatory gene patterns were relevant in human diabetes, we sourced scRNA-seq data from two studies: 1) Bandesh, K. et al 2025^40^ and 2) the human pancreas analysis project (HPAP)^41^, curated by Elgamal, R.M. et al.^92^. 1) The Bandesh, K. et al dataset comprises scRNA-seq data from human Islets of Langerhans, from non-diabetic individuals or individuals with prediabetes or T2DM. A processed seurat object called “Reintegrated transcriptomes of Beta pancreatic islet cells” was available for download at CELLXGENE (https://cellxgene.cziscience.com/collections/58e85c2f-d52e-4c19-8393-b854b84d516e, access: October 4^th^, 2024). 2) The curated HPAP dataset comprises an integrated map of scRNA-seq data from human Islets of Langerhans, from non-diabetic individuals or individuals with T2D or T1D. A processed and annotated seurat object containing this data was downloaded (https://www.gaultonlab.org/pages/Islet_expression_HPAP.html, access: June 6^th^, 2024), and subset to exclusively include β-cells based on the provided annotation “Cell Type Grouped”. These two datasets were used to calculate gene module activity scores of TNF and IFNβ putative target genes in each β-cell, following the procedure outlined in the section “Gene Module Activity”. The TNF and IFNβ target genes were converted from mouse to human gene symbols by mapping mouse gene symbols to Entrez IDs using AnnotationDbi (version: 1.62.2)^89^ and org.Mm.eg.db (version: 3.17.0)^91^. We then matched these mouse Entrez IDs to human orthologs using Orthology.eg.db (version: 3.17.0)^90^ and converted the human Entrez IDs to human gene symbols using org.Hs.eg.db (version: 3.17.)^83^.

### INTERFEROME Database Search

The INTERFEROME database^42^ (v 2.0.1) (analysis run March 6^th^, 2025) is a manually curated database for interferon regulated genes, identified in transcriptomics studies. We submitted the upregulated genes from the high inflammatory subpopulation, identified after three weeks of HFD, to see which of these genes are regulated by interferons and the specific type of IFN it is regulated by. We used the following search conditions: Interferome type: Any, Interferome subtype: Any, Treatments Concentration: Any, Treatment Time: Any, Vivo/Vitro: Any, Species: Mus musculus, System: Any, Organ: Any, Cell: Any, Cell line: Any, Normal/Abnormal: Any, Fold Change Up: 2.0, Fold Change Down: 2.0. Gene symbols were submitted.

### Statistical Analysis

The relevant statistical analysis performed in the single-cell analysis is provided in each section of the section “Single Cell Multiome ATAC + Gene expression sequencing Analysis”. All p-values were adjusted for false discovery rate (FDR) using the Benjamini & Hochberg method when the number of tests exceeded 10. Unless otherwise stated, statistical tests were performed using the nonparametric Wilcoxon test to compare two means. Significance levels were indicated as follows: *: p-value ≤ 0.05. Boxplots depict the first and third quartiles as the lower and upper bounds of the box, with a band inside the box showing the median value; whiskers represent 1.5 times the interquartile range. Point and line plots indicate mean ± SD (standard deviation).

## Notes

https://www.ncbi.nlm.nih.gov/geo/query/acc.cgi?acc=GSE293515

https://github.com/madsen-lab/islets_multiome_phenotyping

https://github.com/madsen-lab/islets_multiome_analysis

### Competing Interest Statement

The authors have declared no competing interest.

### Summary of Updates

Minor revisions. Updated erroneous email addresses and added accession number of sequencing data.

