## Supplementary Figures for "Single Nucleus Multiome Analysis Reveals Early Inflammatory Response to High-Fat Diet in Mouse Pancreatic Islets"

### Supplemental Figure 1

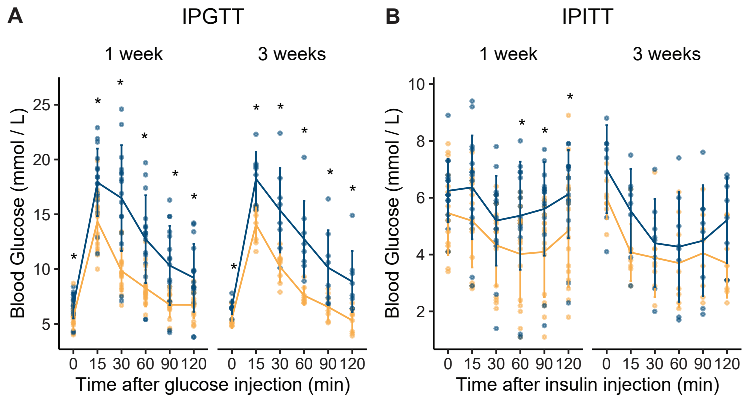

**Supplementary Figure 1. Related to Figure 1.**

11-week-old male C57BL/6JBomTac mice were fed either a low-fat diet (LFD, orange) or a high-fat diet (HFD, blue) for one and three weeks. **A)** Fasted Intraperitoneal Glucose Tolerance Test (GTT) at one week (n = 16 per diet) and three weeks (n = 8 per diet). **B)** Fasted Intraperitoneal Insulin Tolerance Test (ITT) at one week (n = 16 per diet) and three weeks (n = 8 per diet).

### Supplemental Figure 2

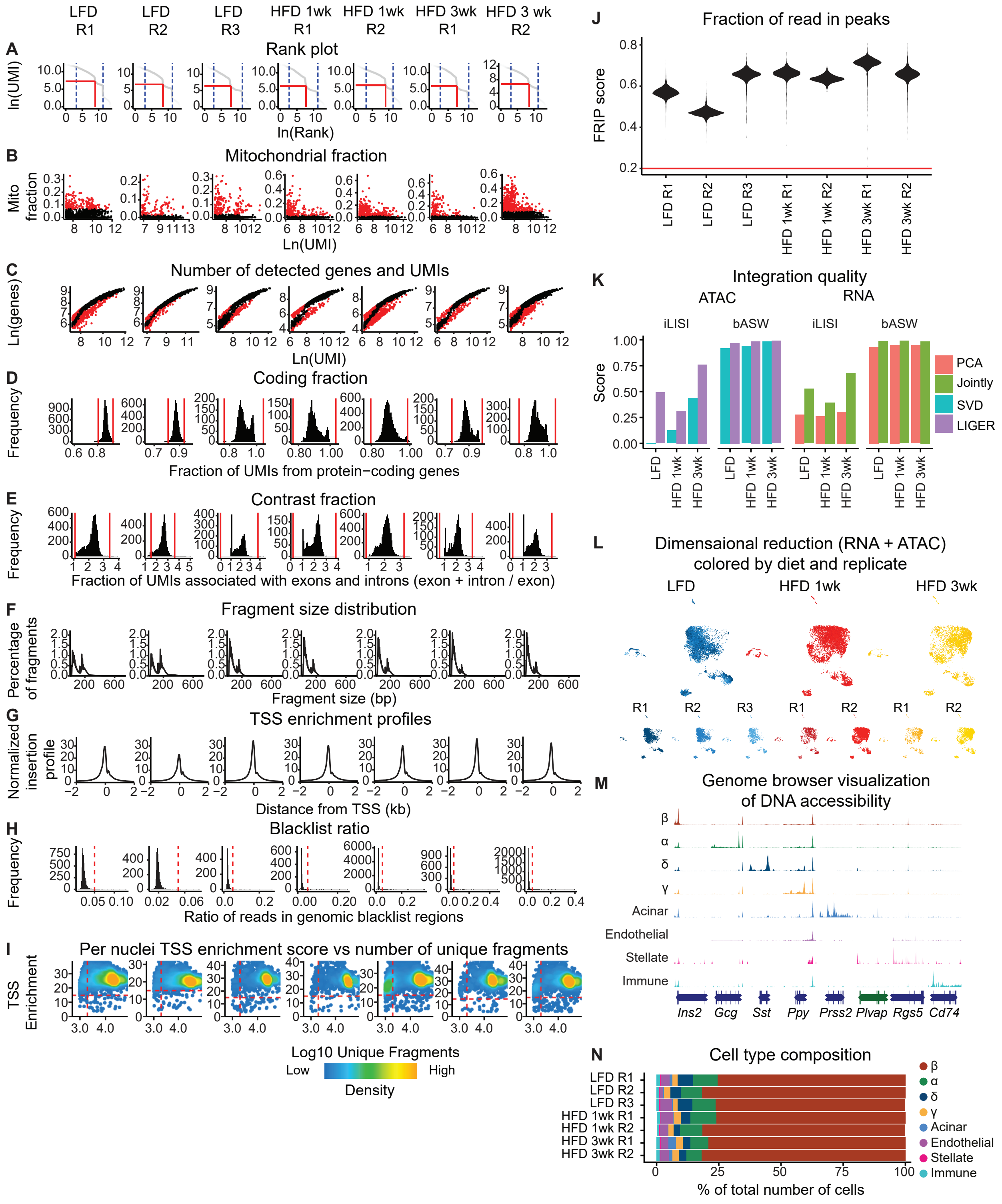

**Supplementary Figure 2. Related to Figure 2.**

**A)** Rank plot of log-transformed total number of unique molecular identifiers (UMIs) (y-axis) and the log-transformed rank in order of decreasing total UMIs (x-axis) for the indicated sample. Vertical red line indicates the rank threshold, and the red horizontal line indicates the corresponding number of UMIs at the threshold. The dashed blue vertical lines represent the quantile range for the breakpoint search, with values between 0.001 and 0.05. These lines define the region within which breakpoints are searched in the barcode ranking distribution. **B)** Scatter plot of the log-transformed number of detected features and the fraction of UMIs associated with mitochondrially encoded genes. Red dots indicate nuclei that did not pass the threshold. **C)** Scatter plot showing the log-transformed total number of UMIs. and the number of detected features. The blue line indicates a piecewise linear regression model with three breakpoints. Red dots indicate nuclei that did not pass the filter. **D)** Histogram showing the distribution of UMIs associated with protein-coding genes. Nuclei falling below or above the red lines were removed. **E)** Histogram showing the distribution of the fraction of UMIs derived from exons (contrast fraction). Nuclei with a contrast fraction below 1 were removed. **F)** Fragment size distribution plot. **G)** Mean Tnf5 insertion frequency around transcription start sites (TSS). **H)** Histogram showing the distribution of the fraction of reads in blacklist regions (blacklist ratio). Nuclei falling above the red line were removed. **I)** Scatter plot showing TSS enrichment score (y-axis) and log10 number of unique fragments (x-axis) for each nucleus. Nuclei falling below the black lines were removed. **J)** Violin plot of the fraction of reads within peaks (FRiP) for each replicate. Nuclei falling below the red line were removed. **K)** Bar plot depicting integration quality based on the ATAC data (left) and RNA data (right). The global integration local Inverse Simpson's Index (iLISI) and the batch average silhouette width (bASW) scores (y-axis) in SVD (blue) -, LIGER (purple) -, PCA (salmon)-, and JOINTLY (green) were calculated for each condition (x-axis). **L)** Uniform Manifold Approximation and Projection (UMAP) visualization of clustering based on a weighted nearest neighbor (WNN) graph, based on the weighted average of RNA and ATAC similarities, split by cluster and replicate. Nuclei are colored according to which condition and replicate they come from as indicated on the figure. **M)** Pseudo-bulk chromatin accessibility tracks scaled between 0-1 for endocrine marker genes by each cell type. Tracks are colored according to cell type as indicated on the figure. **N)** Staged bar plot depicting the percentage (%) of cells in each cell type per condition for each replicate. Bar plots are colored according to cell type as indicated on the figure.

### Supplemental Figure 3

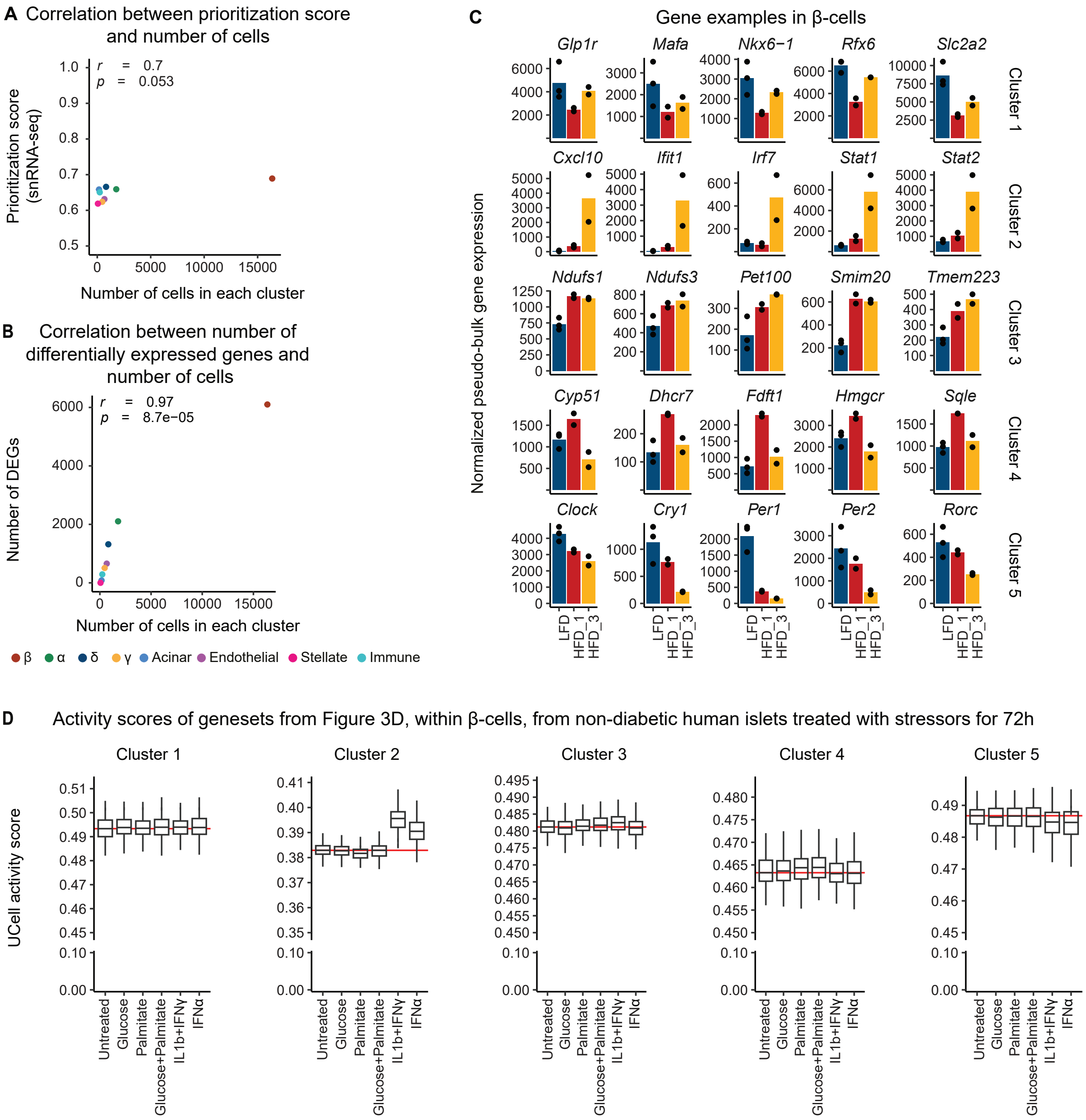

**Supplementary Figure 3. Related to Figure 3.**

**A-B)** Scatter plots depicting the correlation between the number of cells within each cluster and the prioritization score calculated with Augur[21] (A) or the number of differentially expressed genes identified across conditions in each cell cluster (B). Pearson Correlation coefficient (r) and p-value (p) are written inside each plot and points are colored according to cell type as indicated on the figure. **C)** Bar plot of examples of genes from the pathway analysis for each fuzzy cluster found in Figure 3D are shown. Gene expression is depicted for each replicate (points) using DESeq2 normalized pseudo-bulk counts across conditions. Bars indicate the mean expression. **D)** Boxplot of the per cell gene activity score of gene clusters from Figure 3D within pancreatic β-cells from non-diabetic human donors which have been separated into groups treated with stressors; Glucose (22 mM) and/or palmitate (0.5 mM), interleukin-1beta (IL-1β) (1 ng/ml) and interferon gamma (IFNγ) (1000 U/mL), or interferon alpha (IFNα) (2000 U/mL) compared to untreated controls for 72 hours (h) (x-axis)[26,27]. Red lines indicate the median in the untreated samples. Number of donors in each condition at each time point: 72hr; Glucose (n = 2), palmitate (n = 2), glucose and palmitate = (n = 2), IL-1β and IFNγ (n = 2), IFNα (n = 2).

### Supplemental Figure 4

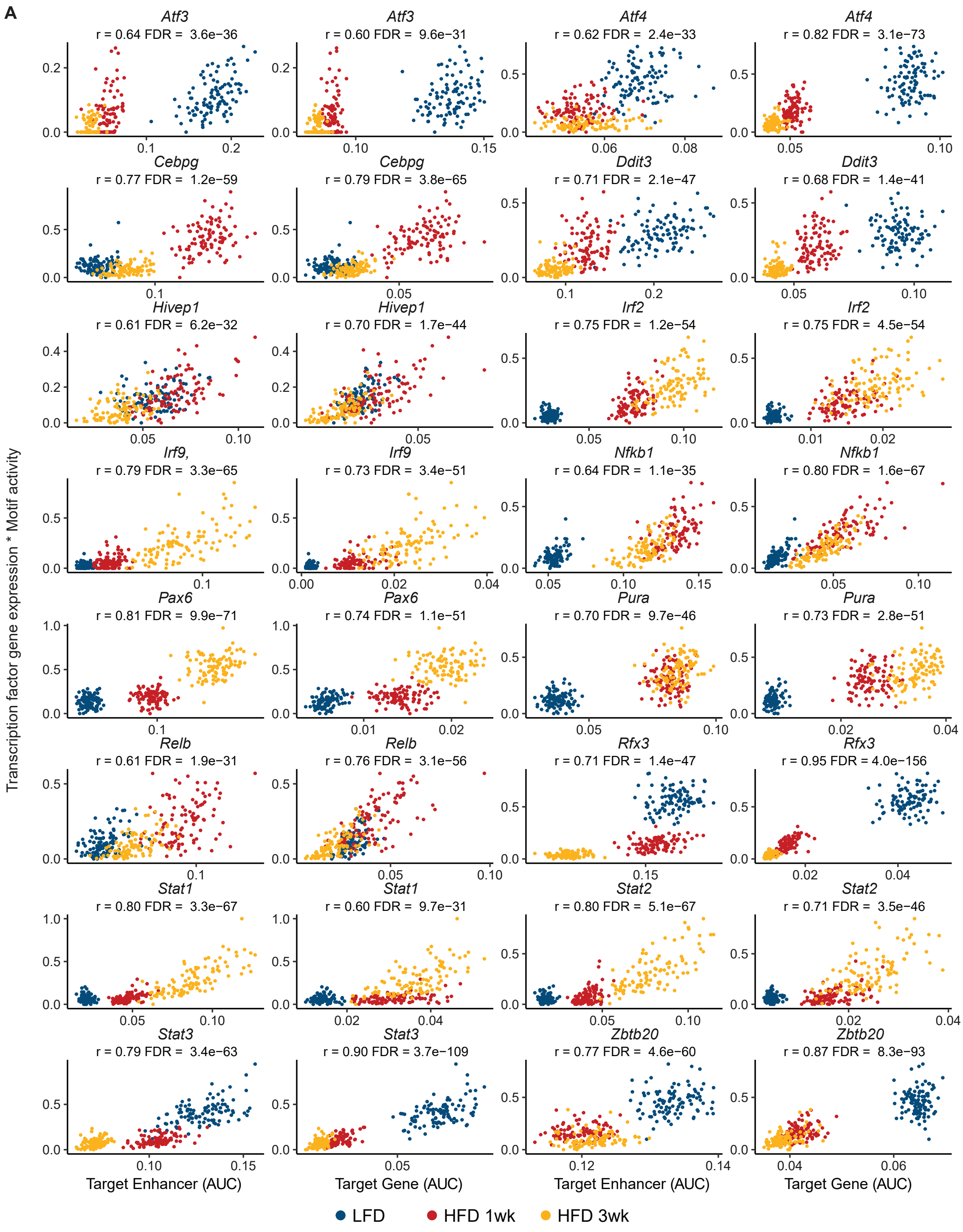

**Supplementary Figure 4. Related to Figure 4.**

**A)** Correlation analysis using semi-pseudo-bulked data (100 meta-cells per condition, each with 10 cells). The correlation analysis was performed using the min-max scaled product of semi-pseudo-bulked transcription factor gene expression and of transcription factor motif activity with activity scores (AUC) for target genes and enhancers from SCENIC+ (computed using AUCell) for each eRegulon (y-axis). Nuclei are colored according to condition as indicated on the figure. Pearson Correlation coefficient (r) is written inside each plot.

### Supplemental Figure 5

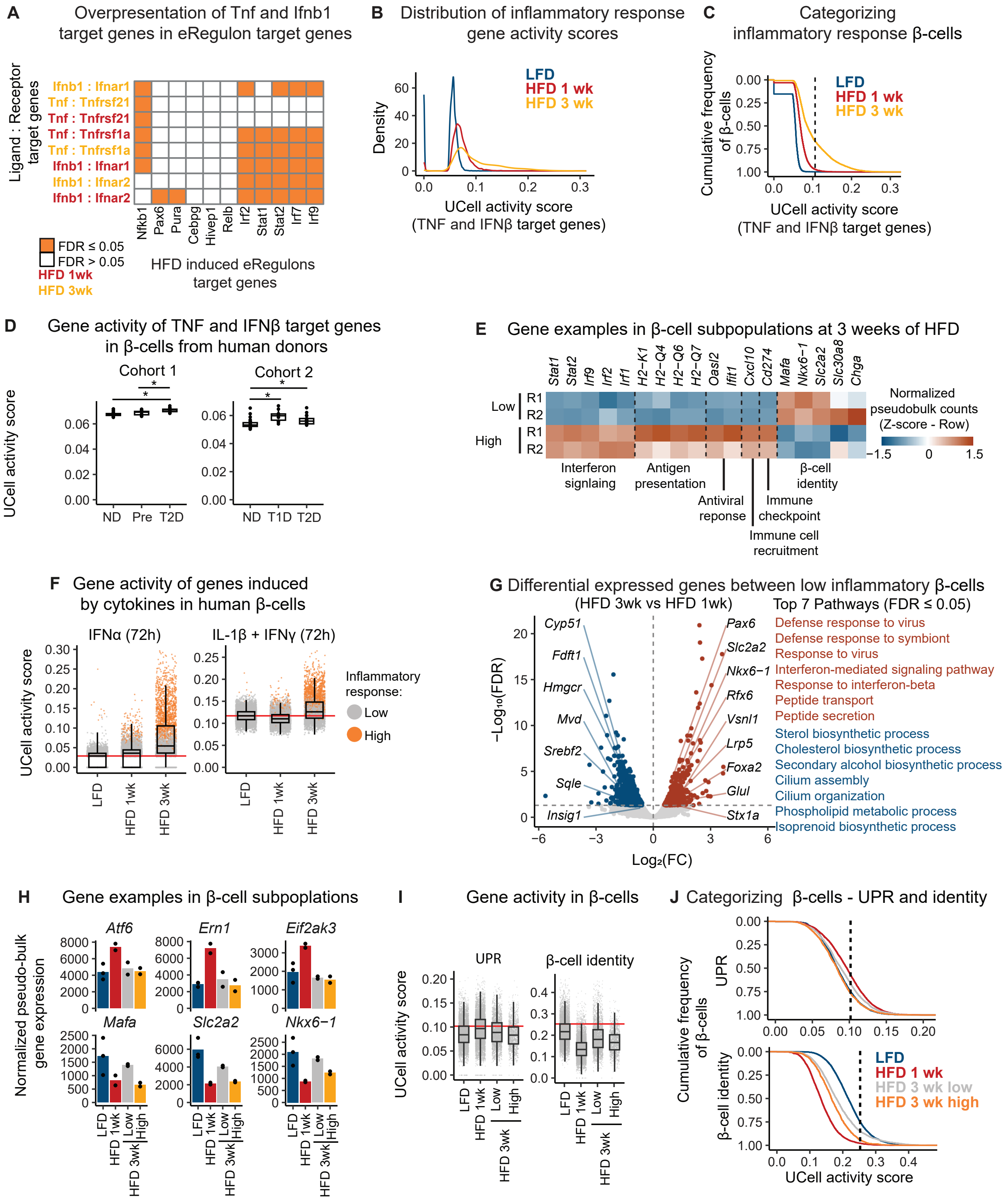

**Supplementary Figure 5. Related to Figure 5.**

**A)** Overrepresentation analysis of target genes from the top TNF and IFN-β interaction partners predicted at one-week HFD (red, HFD 1wk) and three weeks HFD (yellow, HFD 3wk) and eRegulon target genes. Significance was determined using Fisher's exact test. **B)** Density plot showing the distribution of gene activity scores of inflammatory response genes (TNF and IFNβ predicted target genes) split into each condition. **C)** Empirical cumulative distribution function plot of gene activity scores of inflammatory response genes (TNF and IFNβ predicted target genes) split into each condition. The vertical line indicates the threshold used to define cells with high inflammatory signature. **D)** Boxplot of mean gene activity scores using genes for TNF and IFNβ target genes in β-cells from human donors, stratified by donor and disease status in the same way as in Figure 5E. **E)** Heatmap displaying row-scaled z-scores of DESeq2 normalized pseudo-bulk counts for gene examples significantly changed between HFD 3 wk β-cells with low or high inflammatory signature. **F)** Boxplot of the per β-cell gene activity score per condition, using sets of genes that are significantly upregulated in pancreatic β-cells from non-diabetic human donors which have been treated with stressors; IL-1β (1 ng/ml) and IFNγ (1000 U/mL), IFNα (2000 U/mL) compared to untreated controls for 72h[26,27]. Red lines indicate the median in the untreated LFD samples. **G)** Volcano plot depicting differentially expressed genes between β-cell with a low inflammatory response subpopulation at 3 and 1 weeks (3 wk vs 1 wk). The horizontal line indicates false discovery rate (FDR) = 0.05 and the vertical line indicates $\log_2$ fold change (FC) = 0. Selected downregulated (red) and downregulated (blue) genes are annotated. Right panel: The top ten most significant (FDR ≤ 0.05) pathways from Gene Ontology biological processes for upregulated and downregulated genes. **H)** Bar plot of examples of genes related to UPR and β-cell identity. Gene expression is depicted for each replicate (points) using normalized pseudo-bulk counts across conditions, where HFD 3 wk is further divided into low inflammatory (Low) and high inflammatory (High) subpopulations. **I)** Boxplot of gene activity scores using genes related to either UPR (left) or β-cell identity (right), for each condition LFD, HFD 1 wk and HFD 3 wk, where HFD 3 wk samples are subdivided by inflammatory state; low-inflammatory (Low) and high-inflammatory (High). Red line indicates the 75th percentile of scores from LFD. Each point represents a nucleus. **J)** Empirical cumulative distribution function plot of gene activity scores using genes related to either UPR (top) or β-cell identity (bottom) split into each condition. The vertical line indicates the threshold used to define cells with high UPR or identity signature
